# Local chromatin context dictates the genetic determinants of the heterochromatin spreading reaction

**DOI:** 10.1101/2020.05.26.117143

**Authors:** R.A. Greenstein, Henry Ng, Ramon R. Barrales, Catherine Tan, Sigurd Braun, Bassem Al-Sady

## Abstract

Heterochromatin spreading, the expansion of gene-silencing structures from DNA-encoded nucleation sites, occurs in distinct settings. Spreading re-establishes gene-poor constitutive heterochromatin every cell cycle, but also invades gene-rich euchromatin de novo to steer cell fate decisions. How chromatin context, i.e. euchromatic, heterochromatic, or different nucleator types, influences the determinants of this process remains poorly understood. By screening a nuclear function gene deletion library in fission yeast using a previously established heterochromatin spreading sensor system, we identified regulators that positively or negatively alter the propensity of a nucleation site to spread heterochromatin. We find that different chromatin contexts are dependent on unique sets of genes for the regulation of heterochromatin spreading. Further, we find that spreading in constitutive heterochromatin requires Clr6 histone deacetylase complexes containing the Fkh2 transcription factor, while the Clr3 deacetylase is globally required for silencing. Fkh2 acts by recruiting Clr6 to nucleation-distal chromatin sites. Our results segregate the pathways that control lateral heterochromatin spreading from those that instruct DNA-directed assembly in nucleation.

---

Cellular differentiation requires the genome to be partitioned into regions of activity and inactivity such that genes coding for lineage-inappropriate information are repressed. This requires the formation and propagation, in space and time, of gene-repressive heterochromatin structures. Heterochromatin is most commonly seeded by DNA-directed nucleation (Hall et al. 2002; Reyes-Turcu et al. 2011) and then propagates across the chromosome in a DNA-sequence indifferent process termed spreading to repress gene expression in the underlying regions. Silencing is instructed by assembly factors, such as HP1, that recognize heterochromatic chemical modifications (Jacobs et al. 2001; Lachner et al. 2001). The spreading of silencing structures occurs in at least two very different chromatin contexts: (1) Constitutive heterochromatin, which is generally gene-poor and therefore depleted of activities associated with active genes known to antagonize heterochromatin (Scott et al. 2006; Greenstein et al. 2019); or (2) heterochromatin involved in regulating cellular differentiation, which is either seeded at new nucleation sites or invades gene-rich euchromatin de-novo from existing nucleation sites (Wen et al. 2009; Zhu et al. 2013; Zylicz et al. 2015; Zylicz et al. 2018; Nicetto and Zaret 2019). In either scenario, specific factors may intrinsically promote or antagonize this process. For constitutive heterochromatin, maintenance through replication is aided by the inheritance of nucleosomes bearing heterochromatic marks (Alabert et al. 2015). This inheritance promotes modification of nearby nucleosomes due to the “read-write” positive feedback intrinsic to heterochromatin histone modifiers (Zhang et al. 2008; Al-Sady et al. 2013; Ragunathan et al. 2015). When the spreading process invades active chromatin de novo, as occurs in differentiation, it will encounter chromatin modifications that can specifically antagonize heterochromatin (Greenstein et al. 2019). Thus, during this initial invasion process, heterochromatin spreading does not benefit from the inheritance through replication of pre-existing marked nucleosomes. Beyond the differences between active and inactive chromatin, it remains unclear whether distinct nucleation element classes require different regulators to enact efficient spreading outward from those sites.

Fission yeast is an excellent model for addressing the regulation of heterochromatin spreading: (1) It contains a small number of well -defined heterochromatin nucleators, (2) harbors heterochromatin domains that are constitutive as well as others involved in cellular differentiation, and (3) is competent to assemble ectopic heterochromatin domains induced by inserting nucleation sequences into euchromatin. Over the past four decades, forward and reverse genetic screens in fission yeast have established an exhaustive list of factors required for the nucleation of heterochromatin domains. Some of these nucleation mechanisms include repeat sequences that instruct RNAi-machinery to process noncoding (nc) RNAs involved in targeting the histone methyltransferase Clr4 (Moazed 2009); signals within nascent transcripts that trigger heterochromatin island formation (Zofall et al. 2012); pathways involving telomere-protection by the shelterin complex (Wang et al. 2016; Zofall et al. 2016); and transcription factor-bound sequences that recruit heterochromatin regulators directly (Jia et al. 2004). However, less is understood about factors specifically required for the spreading process. With our previously established fluorescent reporter-based heterochromatin spreading sensor, we can segregate the central output of heterochromatin (gene silencing) from the spatial control of the reaction (spreading) (Greenstein et al. 2018; Greenstein et al. 2019). This allows us to address the following questions: (1) Are there known or novel regulators of heterochromatin that primarily regulate spreading, versus nucleation? (2) Does spreading over chromatin with distinct characteristics, such as gene density or nucleosome arrangements, require different sets of regulators? (3) Does the set of regulators required for efficient spreading depend on the type of nucleator that seeds it - for example nucleators using transcription and ncRNA pathways versus direct tethering of heterochromatic factors?

To address these questions, we conducted a series of reverse genetic screens in S. pombe, using a custom collection of nuclear function gene deletions, which we investigated in four different heterochromatin contexts. These include several derivatives of the fission yeast mating type (MAT) locus. This gene-poor constitutive heterochromatin region is contained by IR-L and IR-R boundaries (Grewal and Klar 1997; Thon et al. 2002; Noma et al. 2006) and nucleated by at least two elements: cenH, which is homologous to pericentromeric dh and dg elements and relies on ncRNA pathways, including RNAi (Grewal and Klar 1997; Thon and Friis 1997; Noma et al. 2004); (2) the REIII element, a sequence which recruits heterochromatin factors via the stress-response transcription factors Atf1 and Pcr1 (Jia et al. 2004; Kim et al. 2004). We also queried an ectopic heterochromatin domain that is embedded in gene rich euchromatin. This domain is nucleated by an ectopically inserted dh element fragment proximal to the ura4 locus (Canzio et al. 2013; Marina et al. 2013; Greenstein et al. 2018). The following themes emerge from our genetic screen for heterochromatin spreading in these contexts: (1) The genetic requirements for promotion and containment of heterochromatin spreading differ significantly between distinct chromatin contexts, and to some degree also between different types of nucleators. Among the spreading promoting factors identified, only two contribute to all loci we examined, and neither was previously implicated in heterochromatin assembly. A division of labor between two histone deacetylase (HDAC) complexes in global heterochromatin silencing and spreading. Subcomplexes of the Clr6 complex containing the Fkh2 transcription factor promote spreading, but not nucleation, at multiple heterochromatin loci, while the Clr3 HDAC is required for global heterochromatic silencing at most sites. (3) Broad antagonism of spreading at both euchromatic and heterochromatic loci by a diverse set of nucleosome remodelers, in particular Ino80 and Swr1C.

## Results

Our previously developed heterochromatin spreading sensor relies on three transcriptionally-encoded fluorescent protein-coding genes that collectively report single-cell measurements of heterochromatin formation via flow cytometry, while normalizing for transcriptional and translational noise (Al-Sady et al. 2016; Greenstein et al. 2018). It provides separate, quantitative recordings of nucleation-proximal (“green”) and distal (“orange”) events at a heterochromatin site over large populations of isogenic cells (typically N >20,000, except when strains grow poorly) (**Figure 1A**). In contrast to the singular readout employed by auxotrophy-dependent reporter gene silencing assays, the heterochromatin spreading sensor assay distinguishes changes in heterochromatin nucleation and spreading, and additionally permits tracking of emerging multimodal cell populations and unique population distributions.

**Figure 1:**
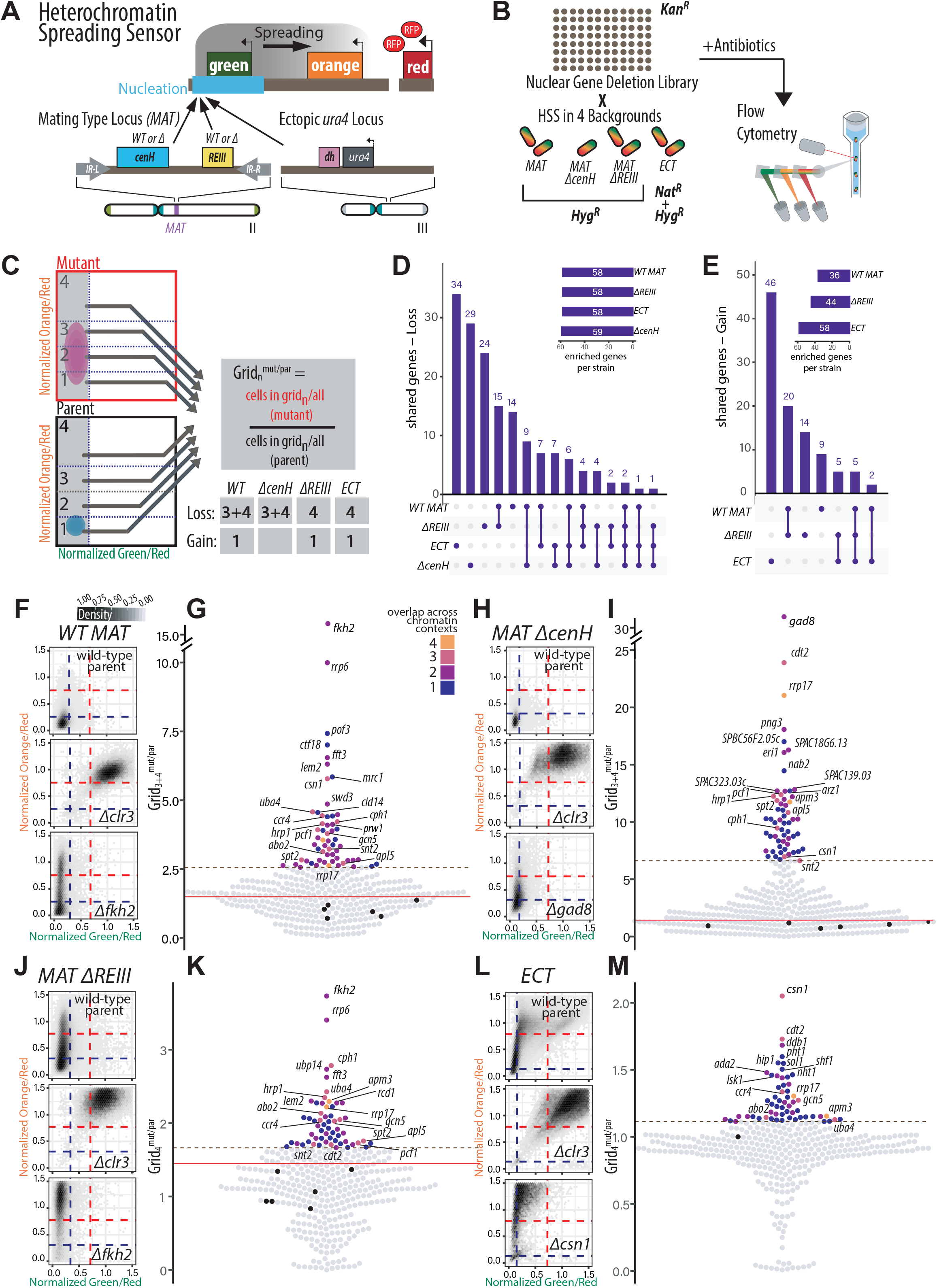
A genetic screen based on a suite of fluorescent reporters identifies heterochromatin spreading regulators in different chromatin contexts. **A.** TOP: Overview of heterochromatin spreading sensor (HSS, Greenstein 2018). Three transcriptionally encoded fluorescent protein genes are integrated in the genome. *SFGFP* (“green”) proximal or internal to the nucleation site allows identification of heterochromatin nucleation; *mKO2* (“orange”) distal to the nucleation site allows identification of heterochromatin spreading. *3xE2C* (“red”) in a euchromatin region normalizes cell-to-cell noise. BOTTOM: The endogenous mating type locus (*MAT*) and ectopically heterochromatic *ura4* locus (Greenstein 2018) were examined with the HSS in the screen. *Bona fide* mutations of the nucleators, *cenH* and *REIII*, in *MAT* were made to limit nucleation to occur from one site. See Figure 1 Supplement 1 for detailed diagrams. **B.** Workflow of the screen to identify genes that contribute to heterochromatin spreading. A custom nuclear function deletion library (Table 1) was mated with four different reporter strains (*WT MAT, MAT ΔcenH, MAT ΔREIII and ECT*). The fluorescence of “green”, “orange” and “red” for each mutant cell within each background are recorded by flow cytometry. **C.** Overview of the spreading-specific analysis with mock distributions of cells and grids indicated. To segregate spreading from nucleation or silencing phenotypes, “green”-off populations (successful nucleation events) are isolated. Within these populations, enrichment of cell populations in particular “orange” fluorescence ranges (Grid_n_) are calculated as Grid_n_. *E.g.* to identify loss of spreading mutants in *WT MAT*, Grid_3+4_ is calculated as percentage of the mutant population divided by percentage of parent population in Grid_3+4_. The Grids used for analysis of gain and loss of spreading in the four chromatin contexts are indicated. **D.-E.** Upset plots indicating the frequency of (**D**) Loss of Spreading, or (**E**) Gain of Spreading gene hits appearing in one or multiple chromatin contexts. For each bar, the chromatin context(s) with shared phenotypes for the underlying gene hits is indicated below the plot. The inset indicates the total number gene hits in each chromatin context of the same phenotype. **F.** *WT-MAT* 2D-density hexbin plots of the wild-type parent, a strong heterochromatin loss hit (*Δclr3*), and the top loss of spreading hit of this chromatin context. Dashed blue lines indicate the values for repressed fluorescence state and dashed red lines indicate values for fully expressed fluorescence state. **G.** Beeswarm plots of Grid_3+4_ for *WT MAT* loss of spreading hits. The top 10 hits are all annotated, and below those hits, mutants that show overlap with at least 3 other chromatin contexts are additionally annotated. Red line, 2SD above the Grid_3+4_ of the wild-type parent isolates (black dots); dashed brown line, the 85^th^ percentile; Dot color, number of chromatin contexts with loss of spreading phenotype over the cutoff. **H.-I.** As in F. and G., but for *WT ΔcenH*. **J.-K.** As in F. and G., but for Grid_4_^mut/par^ of *WT ΔREIII*. **L.-M.** As in F. and G., but for Grid_4_^mut/par^ of *ECT*.

**Table 1:**
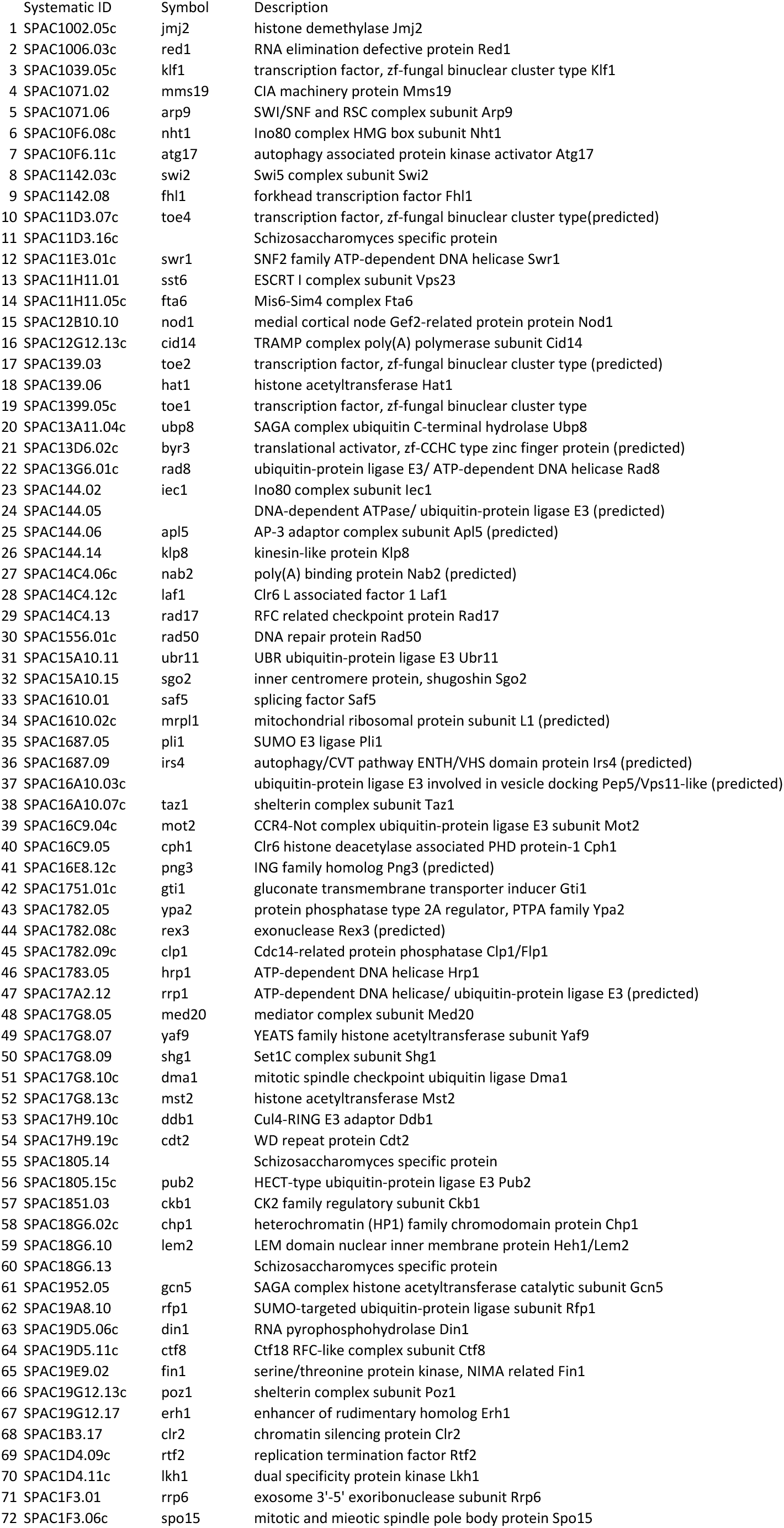
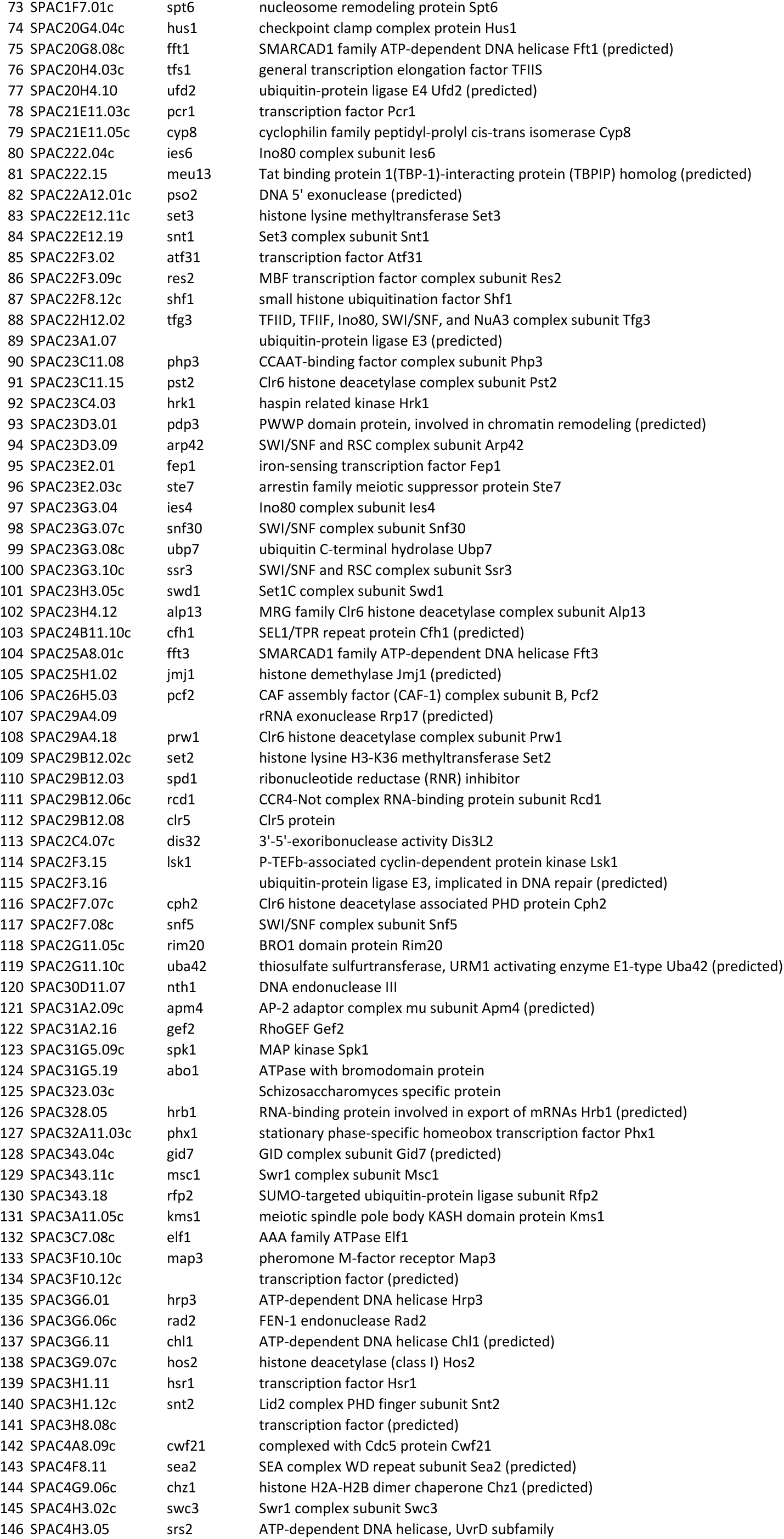
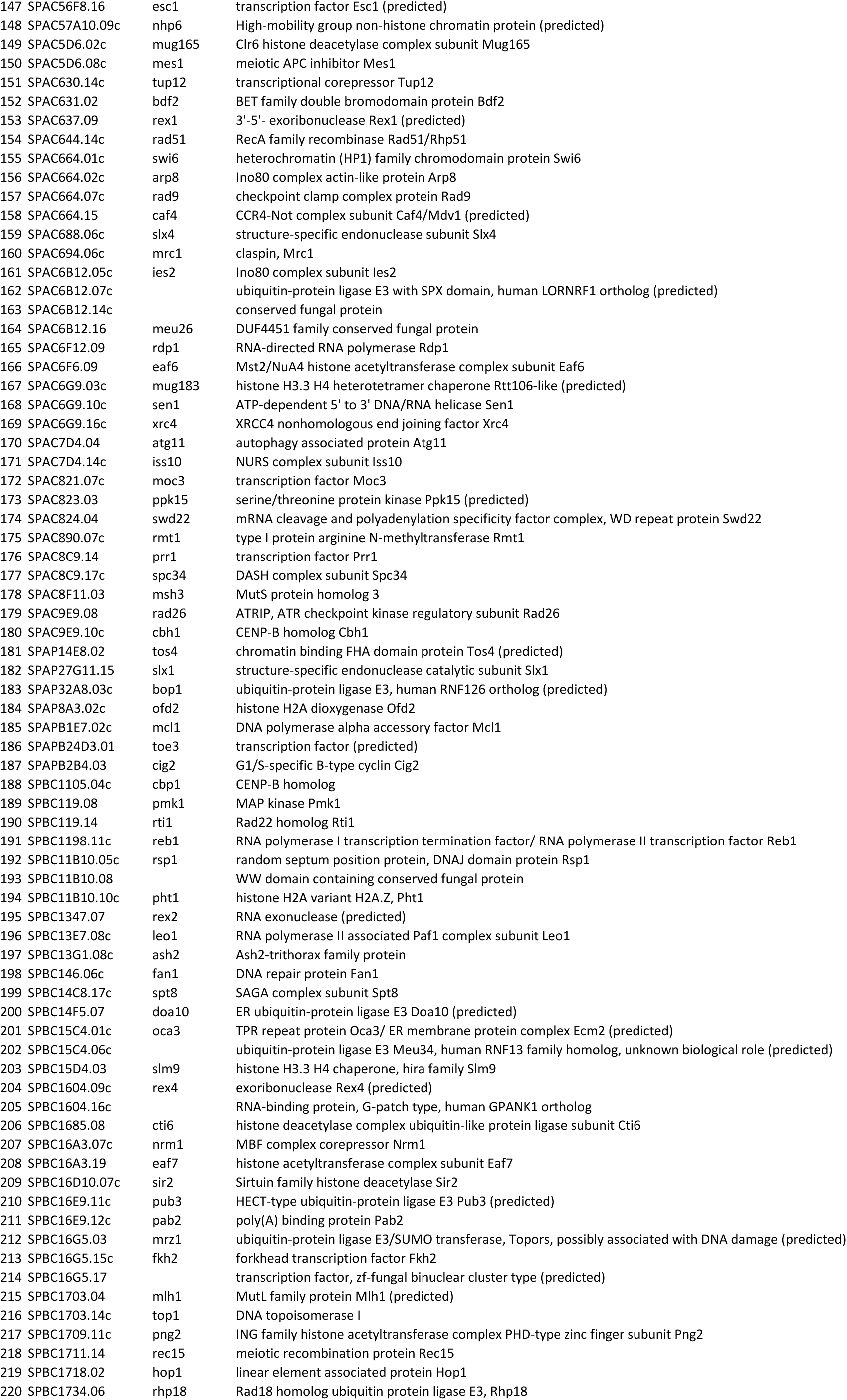
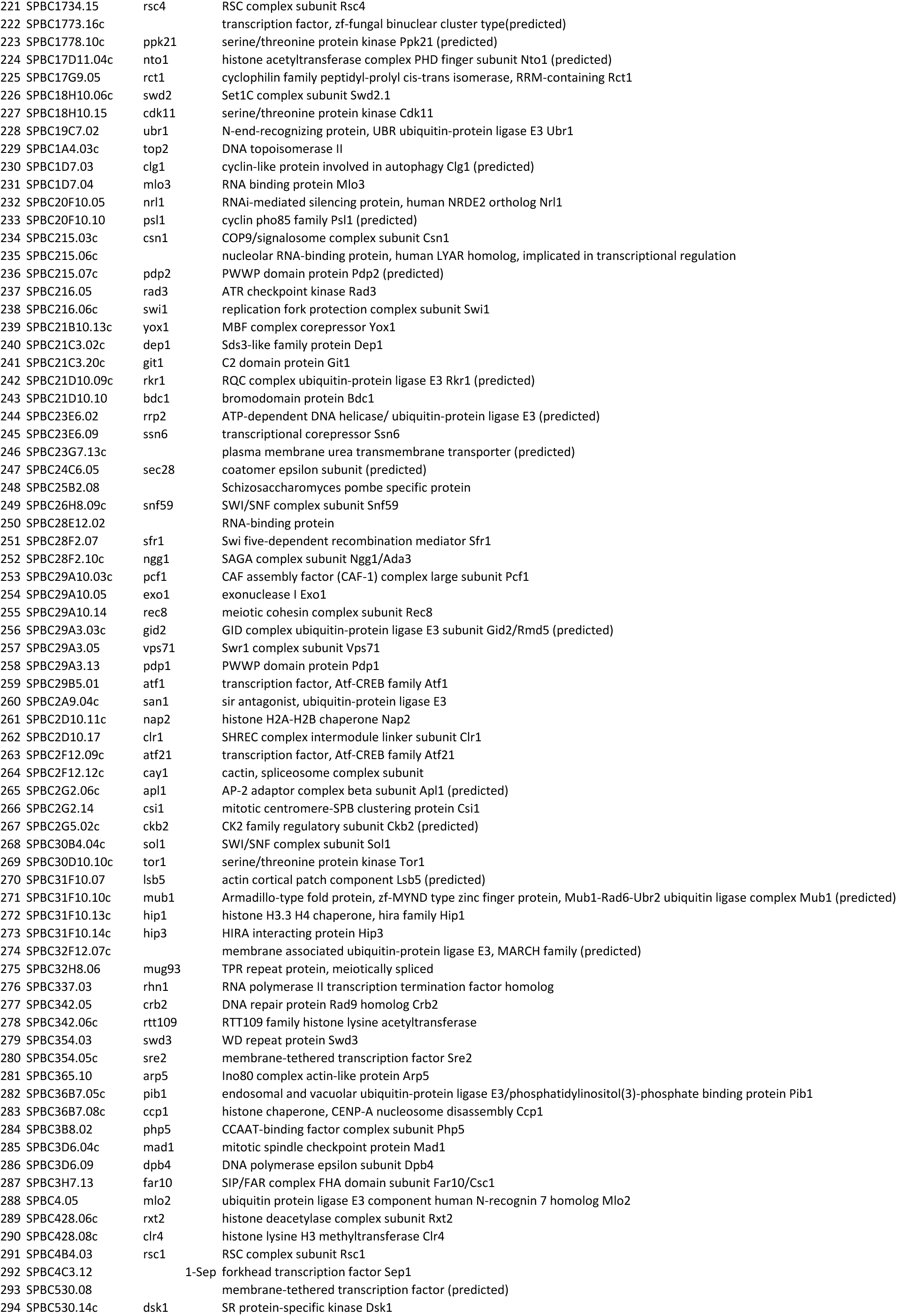
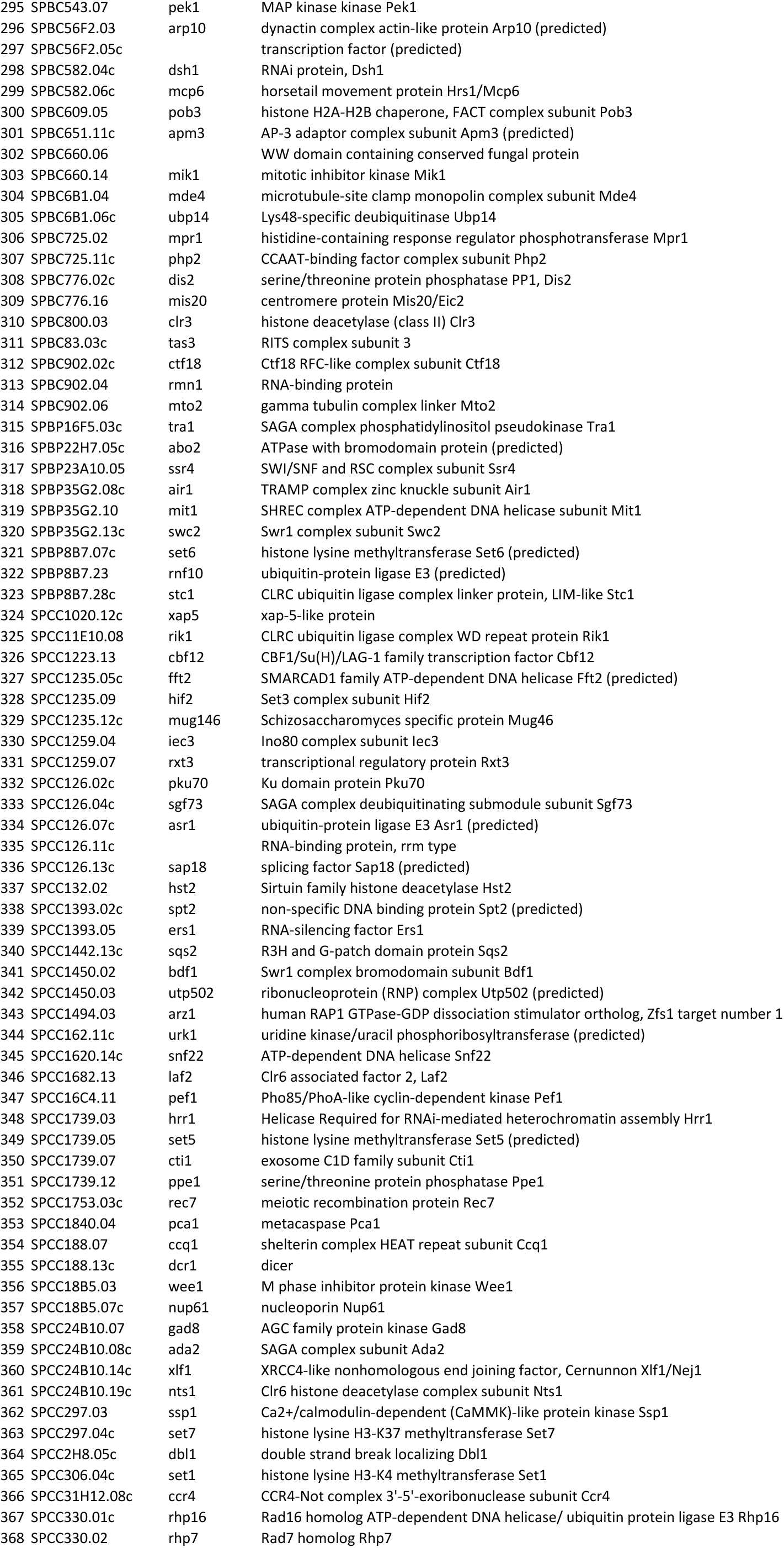
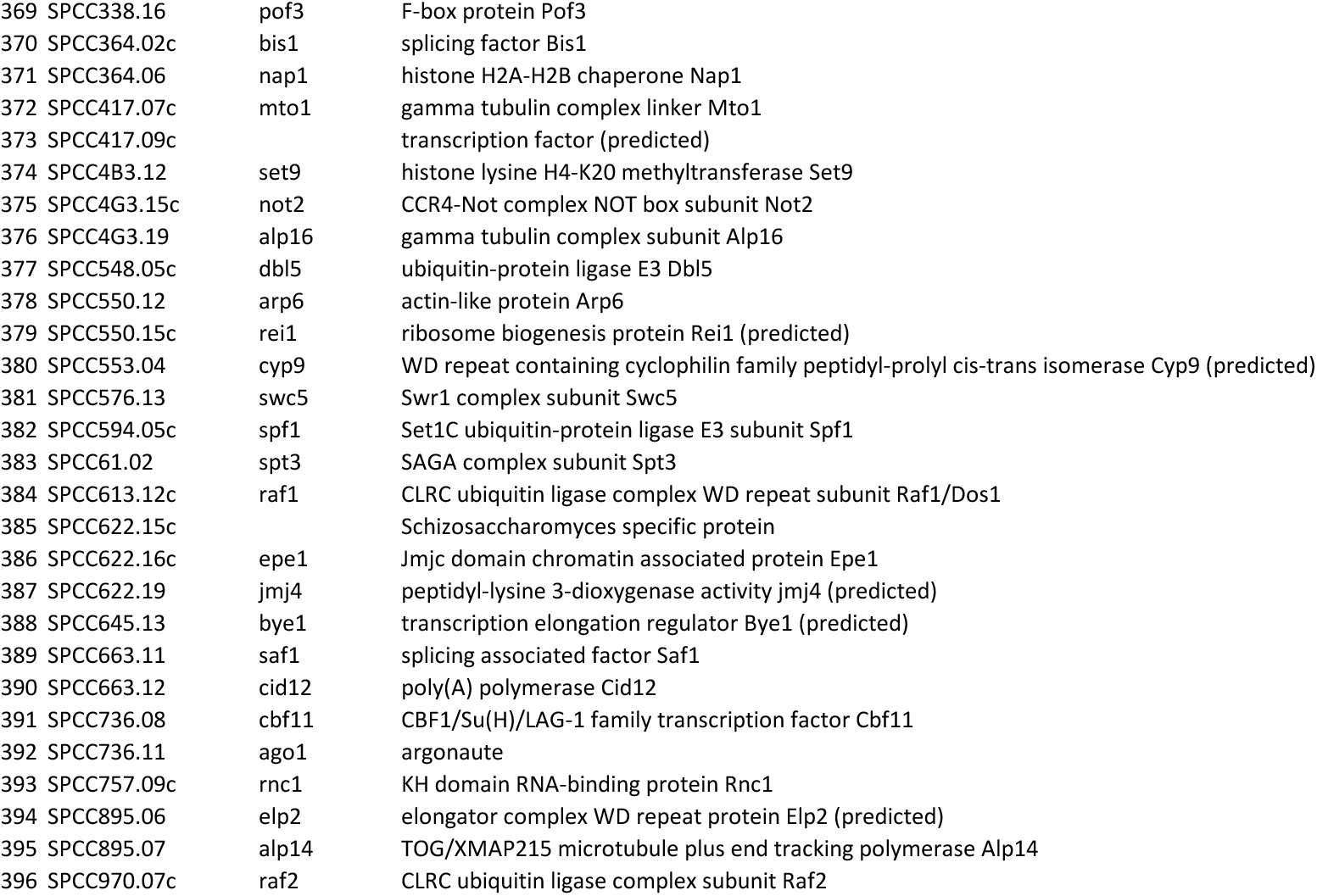
Nuclear function gene deletion library.

**Table 2:**
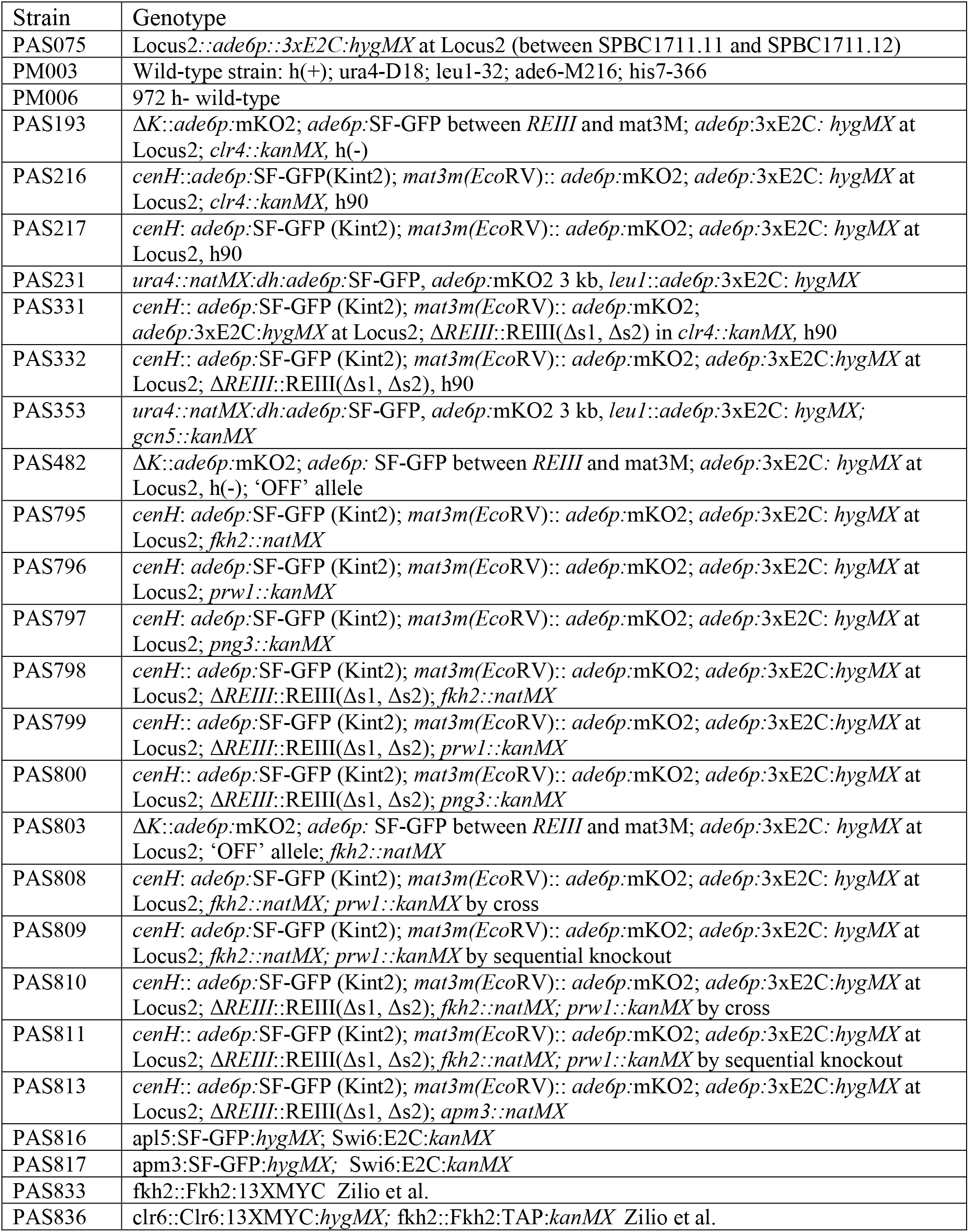
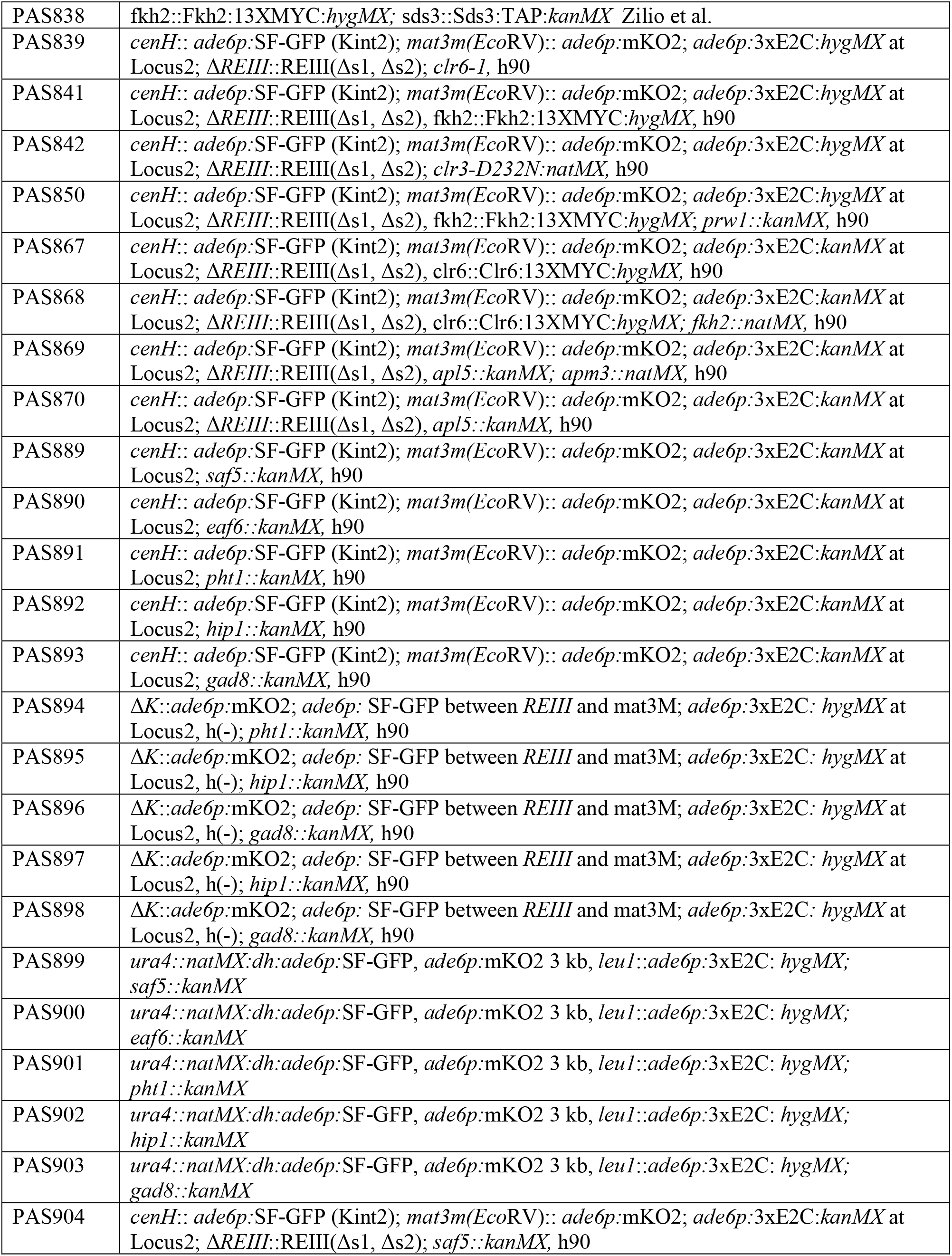
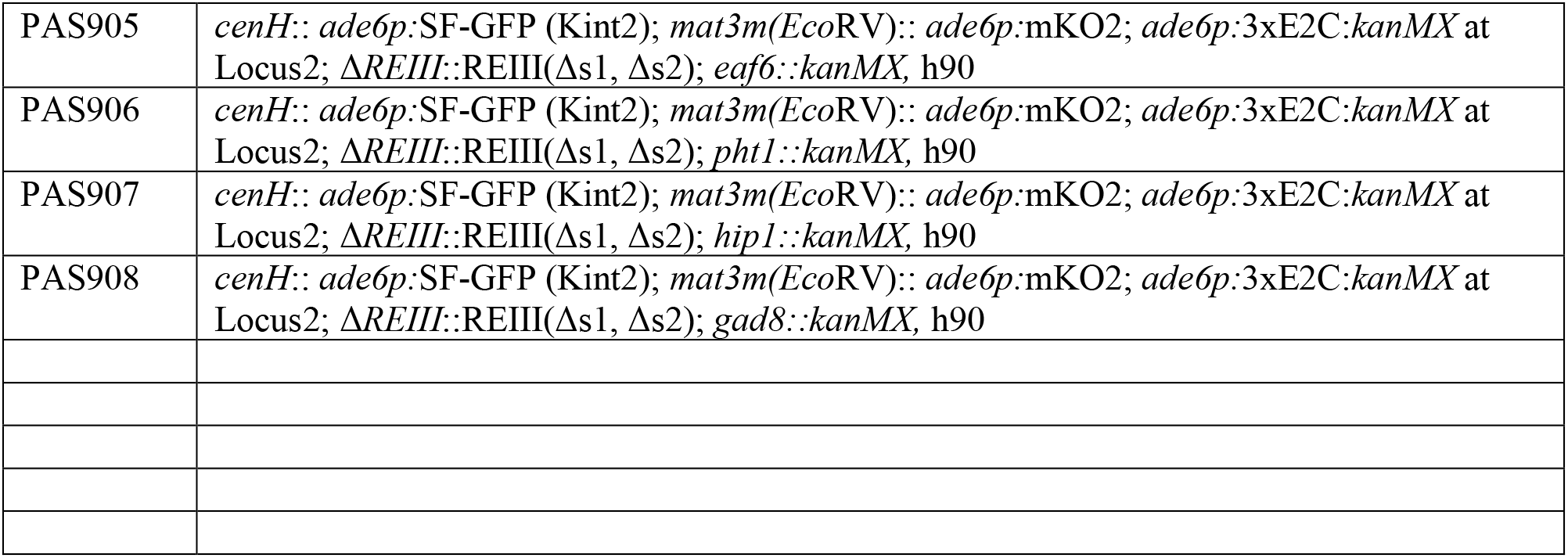
Strain table.

**Table 3:**
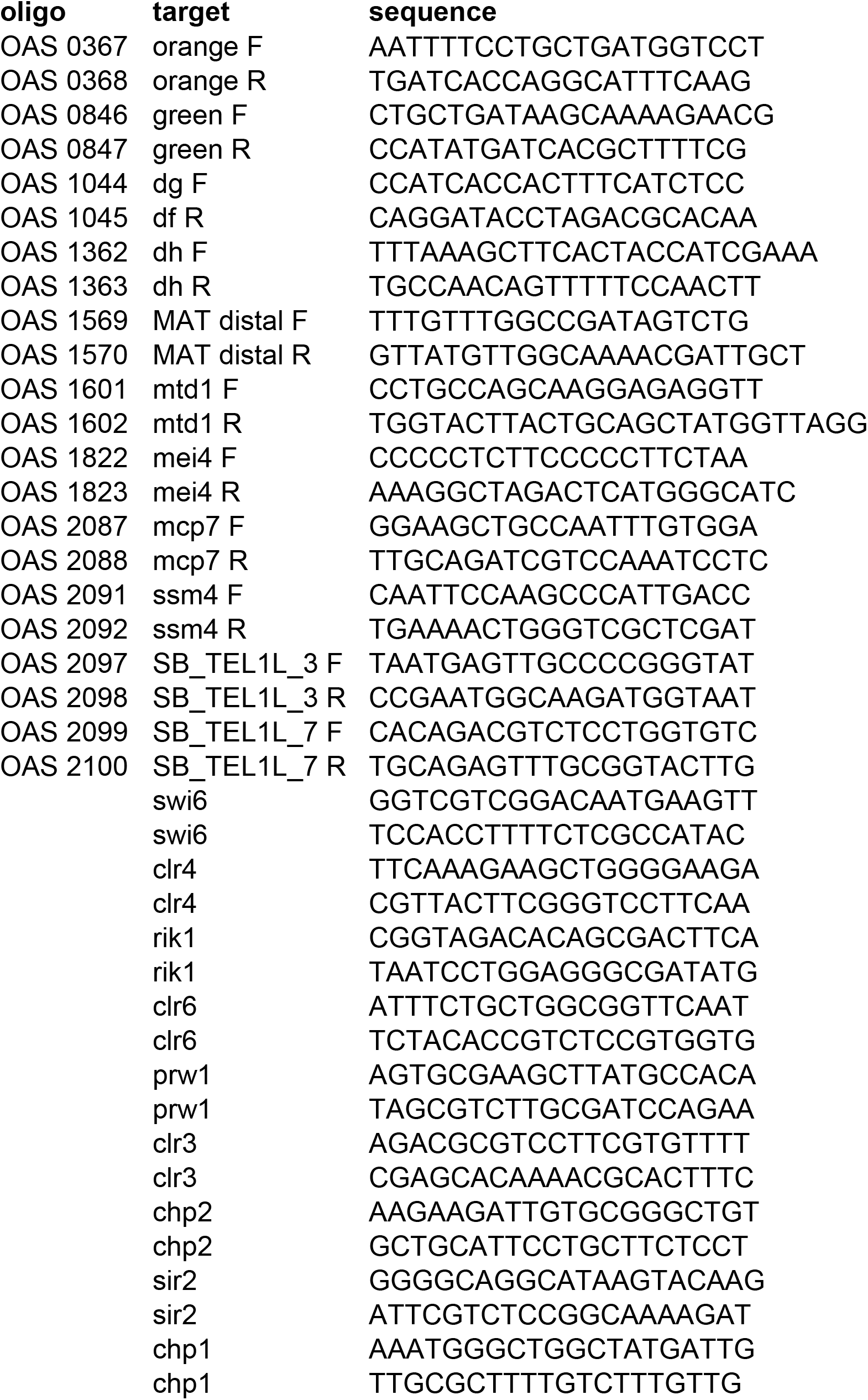
Primers used for ChIP qPCR and RT qPCR.

### Design of heterochromatin spreading sensors at four chromatin contexts

To explore whether the heterochromatin context utilizes general or specific sets of regulators for the spreading reaction, we queried three different derivatives of the constitutively heterochromatic mating type (MAT) locus and one euchromatic context, each containing an embedded heterochromatin spreading sensor (**Figure 1A, Figure 1 Supplement 1**) (Greenstein et al. 2018). The mating-type locus contexts included wild type MAT, with the cenH and REIII nucleating DNA elements intact, and two MAT variants that contained mutations in either the cenH or REIII elements (**Figure 1A, Figure 1 Supplement 1A,B**). Mutations in these DNA elements limit initiation of heterochromatin spreading to one nucleator (Greenstein 2018). To probe heterochromatin formation in the euchromatic context, we focused on the ura4 locus, where heterochromatin spreading is ectopically driven by the upstream insertion of a pericentromeric dh DNA element (**Figure 1 Supplement 1C**, (Marina et al. 2013; Greenstein et al. 2018)). We refer to this chromatin context as ECT (ectopic). When analyzed by flow cytometry, WT MAT and MAT ΔcenH (OFF isolate, see methods) populations appear fully nucleated with near-complete local spreading, as evidenced by population density in the bottom left in the 2D density hexbin plot (**Figure 1F,H TOP** (Greenstein et al. 2018)). MAT ΔREIII and ECT cell populations, while mostly nucleated, display a stochastic distribution of spreading states, as evidenced by a vertical distribution on the left of the 2D density histogram (**Figure 1J,L TOP** (Greenstein et al. 2018)). While the distance between nucleation and sensor sites varies slightly for the different chromatin context analyzed (from 2.4 to 3.6 kb; see **Figure 1 Supplement 1**), we showed previously that altering the distance between “green” and “orange” does not qualitatively affect the output (Greenstein et al. 2018). Thus, we presume that these differences in behavior reflect different intrinsic properties of the chromatin environment rather than differences in the reporter arrangement. In addition to wild-type background, we display Δclr3 as reference for a strong loss-of-silencing mutant (**Figure 1F,H,J,L**).

### Identification of chromatin context-specific spreading regulators

We crossed a deletion library of ∼400 nuclear function genes (see methods and Table 1, Figure 1B) to these four reporter strains and measured nucleation and spreading by flow cytometry. To segregate spreading from nucleation, we first isolated cell populations that reside within a “green”- off gate, which represents cells with heterochromatin fully assembled at the nucleation site and no expression of the reporter (see methods, Figure 1C and (Greenstein et al. 2018)). Within this gate, we divide the “orange” signal into 4 grids, from fully repressed (i.e., complete spreading over “orange”; Grid 1) to fully de-repressed (i.e., no silencing at “orange; Grid 4), with the remaining space symmetrically divided to yield Grids 2&3 (see methods). To quantify increased or decreased spreading in a given mutant, we calculated a Gridn^mut/par^ metric (described in methods), which tracks the changes of cell distributions in “orange” expression within the “green”-off gate (**Figure 1C**). Since WT MAT and MAT ΔcenH display very tight silencing of both “green” and “orange” with very few events in grid 4 (**Figure 1F,H**), we used a Gridn^mut/par^ metric that considers both grids 3+4 for the robust identification of spreading defects. MAT ΔREIII and ECT have a more stochastic spreading behavior with “green”-off cells populating a range of “orange” states from OFF to ON (Greenstein et al. 2018), including to some extent both grid 3 and 4 (**Figure 1J.L**). Hence, for these two contexts we used a Gridn^mut/par^ metric that only considers grid 4 to focus on complete loss of spreading (i.e. “orange” signal in the range of Δclr4 control). To isolate gain or loss of spreading mutants (hits) for further analysis, we employed two metrics (1) a significance threshold when multiple parental isolates were available (all except ECT) and (2) a percentile based cut-off. As significance threshold, we only considered mutants in which Gridn^mut/par^ values were above 2 standard deviations beyond the mean of the parent isolates (Figure 1 G,I,K,M). As an additional cut-off, we only considered the top 15% of all Gridn^mut/par^ -ranked mutants, even if more genes pass the 2SD significance threshold. Having identified these gene hits, we next proceeded to analyze their relationships within and across chromatin contexts.

We first examined the degree to which spreading modulators are shared between chromatin contexts via upset plots (**Figure 1D,E**). Conceptually similar to a Venn diagram, this analysis allows rapid visualization of the degree of overlap between sets, with the number of shared hits plotted as a bar graph and the sets each bar represents annotated below the plot. For loss of spreading phenotypes (i.e. genes that promote spreading, Figure 1D), these upset plots showed that exceedingly few genes (i.e. 2 of 164 unique hits) are shared across all chromatin contexts, emphasizing the specific impact of each context on heterochromatin spreading. Notably, these two genes (*apm3, rrp17*) have not previously been implicated in heterochromatin assembly. Apm3 has been proposed to be part of an AP-3 adaptor complex, whereas Rrp17 is a predicted rRNA exonuclease. Six genes were shared across all the MAT locus chromatin contexts (*apl5, cph1, hrp1, spt2, snt2, pcf1*). In contrast, the majority of hits (i.e. 101 genes) contributed only to one chromatin context. The degree to which genes contributed positively towards spreading (Gridn^mut/par^) and the degree of overlap across chromatin contexts (by color code) is shown in **Figure 1G,I,K,M**. We additionally show the 2D density hexbin plot for the top loss-of-spreading hit for each context (**Figure 1F,H,J,L**). Among the top hits, we made the following observations: (1) Pathways involved in the removal of the heterochromatin antagonist Epe1 play a prominent role in promoting spreading. Epe1 is turned over by an E3 ligase complex (Braun et al. 2011), which includes the adaptor proteins Ddb1 and Cdt2 and is regulated by the COP9 signalosome also involved in Epe1 turnover (Bayne et al. 2014).We found that *csn1* (COP9), *cdt2*, and *ddb1* are the three strongest hits in ECT and that cdt2 is also among the top hits in MAT ΔcenH. (2) Beyond the Epe1 turnover pathway, additional top hits unique to ECT are the histone variant H2A.Z (*pht1*), which antagonizes heterochromatin spreading in budding yeast (Meneghini et al. 2003), and an Asf/HIRA chaperone subunit (*hip1*) (Figure 1M). (3) For WT MAT and MAT ΔREIII, we found as top hit fkh2, which encodes a transcription factor, and as second top hit, *rrp6*, a key member of the nuclear exosome (see below; **Figure 1G,K**). (4) The strongest and largely unique hit in MAT ΔcenH is gad8, which encodes a protein kinase that targets several factors, including Tor2 and Fkh2 (**Figure 1I**).

As mentioned above, loss of the AP-3 adaptor complex subunits Apm3 and Apl5 induced loss of spreading in either all chromatin contexts (Apm3) or only the MAT contexts, respectively (Apl5, also observed in (Holla et al. 2020)). We further assessed the activities of Apm3 and Apl5 by newly generating *Δapm3* and *Δapl5* single and double mutants in the MAT ΔREIII background, where *Δapm3* and *Δapl5* had a moderate and mild effect in the screen, respectively. We reproduced the mild to moderate spreading defect for both single mutants; we further observed a slightly stronger defect in the *Δapm3 Δapl5* double mutant (Grid ^mut/par^ 1.56) compared to the *Δapm3* and Δapl5 single mutants (Grid ^mut/par^ 1.4 and 1.16, respectively); **Figure 1 Supplement 2A-D**). Whereas Apl5 is largely cytoplasmic, Apm3 shows both nuclear and cytoplasmic localization, (**Figure 1 Supplement 2E,F**) and also affects H3K9me2 accumulation at heterochromatin islands (**Figure 1 Supplement 2G**). Together, these findings may support the notion of a direct rather than indirect role for Apm3 in heterochromatin assembly. However, further work is needed to elucidate the specific function of Apm3 in heterochromatin spreading and whether this is linked to the AP-3 complex itself.

Besides loss of spreading, we also identified mutants that showed gain of spreading in WT MAT, MAT ΔREIII and ECT (Figure 1E, Figure 1 Supplement 3) by examining a Gridn^mut/par^ metric that considers grid 1. We could not examine MAT ΔcenH for this phenotype because this chromatin context is highly repressed in the OFF state as reported previously (Grewal and Klar 1996; Greenstein et al. 2018). Also for this phenotype, only a limited number of mutants are shared as hits across these three chromatin contexts (5 out of 98, *vps71, arp6, leo1, git1, pmk1*, Figure 1E; examples for top hits are shown in Figure 1 Supplement 3). Notably, Leo1 was previously shown to be implicated in spreading control across boundaries (Verrier et al. 2015), whereas Vps71 and Arp6 are members of the H2A-Z-specific Swr1 remodeling complex(Krogan et al. 2003). ECT displayed the largest fraction of spreading-antagonizing genes that are unique to a single context, with 46 out of 69. In contrast, while ECT displays a very similar spatiotemporal spreading behavior to MAT ΔREIII (Greenstein et al. 2018), we found little overlap between the two and only 14 genes unique to MAT ΔREIII (**Figure 1J,L**). The larger number of antagonists in ECT suggests that spreading is under additional layers of control in the euchromatic context.

Due to the chromatin context-specific effects and the fact that this type of spreading analysis with the heterochromatin spreading sensor system has not been conducted before, we sought to independently validate above observations. We selected 5 mutants that have chromatin context-specific effects, covering both loss and gain of spreading: *saf5*, which shows gain of spreading in WT MAT and mildly in MAT ΔREIII; *eaf6*, which shows gain of spreading in MAT ΔREIII only; *pht1* and *hip1*, which show loss of spreading only in ECT; and *gad8*, which shows loss of spreading primarily in MAT ΔcenH. We recreated these mutations de novo in the chromatin contexts described above and conducted RT-qPCR analysis for SF-GFP (“green”, nucleation) and mKO2 (“orange”, spreading) transcripts. This validation approach recapitulated broadly what our screen analysis showed **(Figure 1, Supplement 4**), confirming the context-specific functions in heterochromatin spreading of these gene products.

Taking our analysis beyond individual genes, we sought to query which protein complexes are involved in spreading regulation, as this can point out the major pathways involved in this process. Using the Gene Ontology (GO) protein complex annotations from Pombase (Lock et al. 2018), we annotated each gene hit identified using the criteria described in Figure 1 (G-M). We then tabulated the frequency (“counts”) of each GO complex per chromatin context for both loss of spreading (loss) and gain of spreading (gain) phenotypes, performed unsupervised clustering on the data, and displayed the results as a heatmap (**Figure 2 Supplement 1**). Overall, we identified three major common trends: (1) a role for chromatin remodelers in antagonizing spreading; (2) a role for the SAGA complex in promoting spreading at ECT; (3) a role for Clr6 histone deacetylase complexes (HDACs) in promoting spreading, with the notable exception of Set3C, which antagonizes spreading and is annotated into the expanded Rpd3L complex.

### Chromatin remodelers broadly antagonize spreading

Mutants defective in chromatin remodeling complexes strongly contribute to the “gain” phenotype category, including the Swr1C, Ino80, SWI/SNF, and RSC-type complexes (**Figure 2 Supplement 1**). To explore this further, we assessed which protein components contributed to these GO complex counts. For all genes annotated to a given chromatin remodeling complex and present in our screens, we displayed whether they were identified as a hit (blue) or not (grey) in a hit table (**Figure 2A**). Indeed, we found that the large majority of the gene hits annotated fall within the “gain” but not “loss” phenotype across backgrounds, confirming a potential role of these nucleosome remodeling complexes in antagonizing spreading.

**Figure 2:**
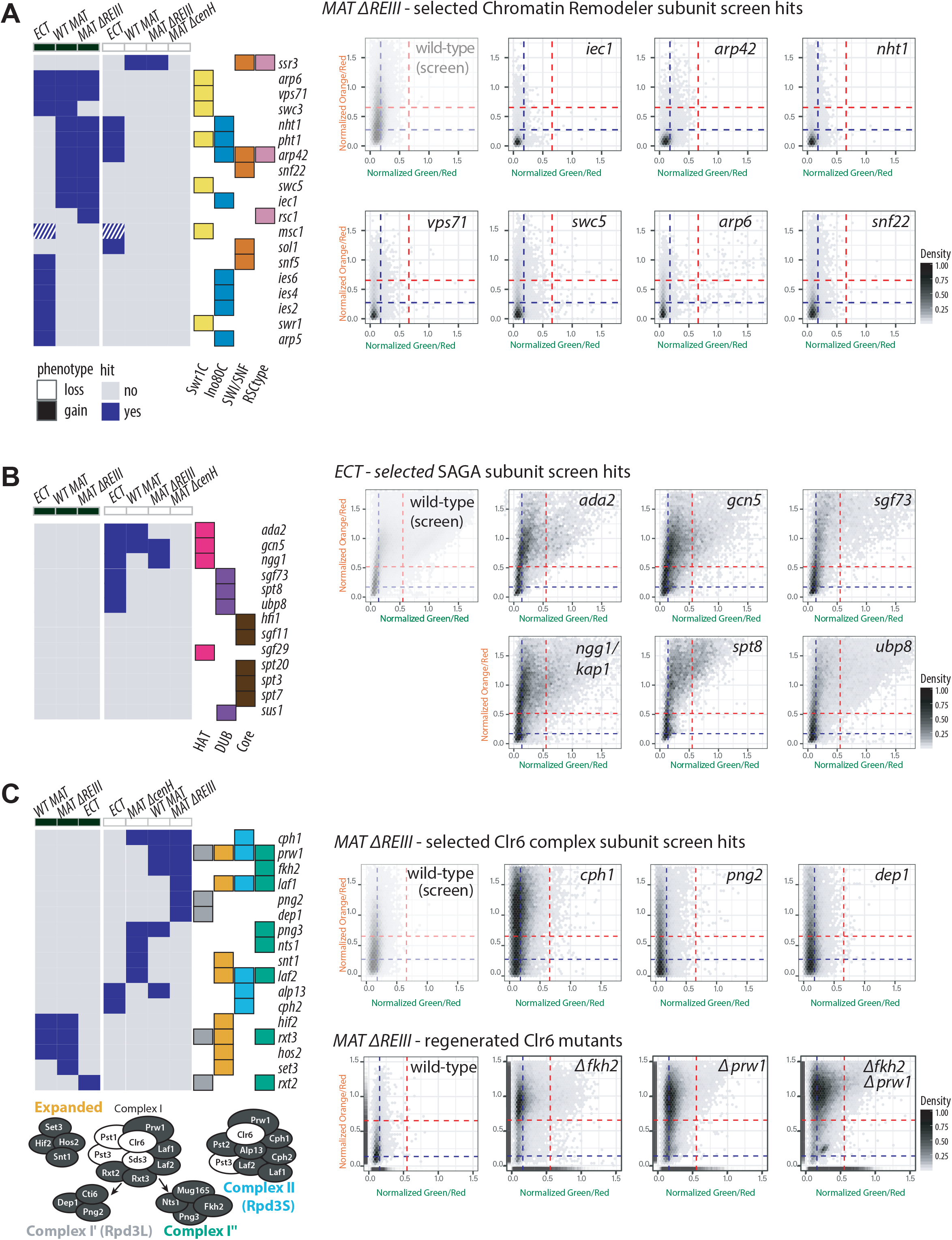
Chromatin remodeler, SAGA and Clr6 complexes regulate heterochromatin spreading. LEFT: Table of complex members which were hits: **(A)** Swr1C, Ino80, SWI/SNF, and RSC-type chromatin remodeler complexes, **(B)** the SAGA complex, and **(C)** The Clr6 complexes Rpd3L, Rpd3L-Expanded, Rpd3S, and Clr6 Iʺ (Bottom shows schematic of Clr6 complexes, subunits indicated in white are essential). Components identified as a hit in either showing a “gain” or “loss” of spreading phenotype for each background are marked blue, subunits that are a hit for both phenotypes (*msc1*) as white-blue crosshatched. The proteins present in each complex or subcomplex are annotated at the right with the presence of color indicating membership of that protein in the complex. For remodelers and Clr6 complexes, subunits that are not hits in any chromatin context are not included, for SAGA, all subunits except TAF_II_s are shown. For **(C)**, Clr6 mutants that were not hits are *mug165, pst2* and *cti6*. RIGHT: 2D density hexbin plots for selected screen hits from different families of complexes in respective background: **(A)** Chromatin remodeler gain of spreading screen hits for *MAT ΔREIII,* (**B**) SAGA loss of spreading screen hits for *ECT*, and **(C)** Clr6 complex subunit loss of spreading screen hits in *MAT ΔREIII, as well as de novo* generated Clr6 complex subunit deletion mutants. *MAT ΔREIII* and *ECT* parents shown in Figure 1 are shown here again (with transparency) for comparison. *Δfkh2, Δprw1 and Δfkh2 Δprw1* mutants were re-created *de novo* in *MAT ΔREIII.* A rug plot is included on the X and Y axes indicating the 1D density for each color. Rug lines are colored with partial transparency to assist with visualization of density changes.

### SAGA promotes spreading, primarily in the euchromatic context

A surprising observation was the enrichment of a large number of subunits of the SAGA complex among the loss-of-spreading hits, which was represented with six subunits in the ECT context, the most enriched single complex for any background (**Figure 2B**). This implies that SAGA, a histone acetylase involved in gene transcription, positively regulates heterochromatin spreading in euchromatin. To assess if SAGA contributes directly to spreading of the heterochromatin structure, rather than indirectly to gene silencing, we assessed the accumulation of H3K9me2 at “green” and “orange” in the ECT background, as well as the pericentromeric *dg* element. Consistent with the screen data, we find that the histone acetylation catalytic subunit Gcn5 is required for efficient spreading of H3K9me2 to “orange”, but not its establishment at “green” or at *dg* **(Figure 2, Supplement 2**).

### Clr6 HDAC complexes promote spreading, primarily in constitutive heterochromatin

Three classes of HDACs exist, which have partially redundant and non-overlapping functions in the formation of heterochromatin domains and gene silencing. Clr6 belongs to class I and is part of several sub-complexes, contributing to both heterochromatic and euchromatic gene regulation (Grewal et al. 1998; Nicolas et al. 2007). Clr3 belongs to class II and is a member of the SHREC complex (Sugiyama et al. 2007) and Sir2 belongs to the class III HDAC of the sirtuin family (Shankaranarayana et al. 2003). Unlike class II and III HDACs (Clr3, **Figure 1**; Sir2, **Figure 1 Supplement 5**), we show here that sub-complexes of the Clr6 family, including the Rpd3S, Rpd3L, and Clr6 Iʺ, appear to contribute exclusively to spreading but not nucleation (for hit table of top hits and 2D hexbin plots see Figure 2C; for additional 2D hexbin plots of all Clr6 components see **Figure 2 Supplement 3**). As noted above, the forkhead transcription factor Fkh2 was identified among the strongest spreading hit in WT MAT and MAT ΔREIII HSS strains. Despite not being formally annotated to the Clr6 Iʺ complex by GO terms, Fkh2 was previously described as a member of this sub-complex (Zilio et al. 2014). We included Fkh2 as a member of Clr6 Iʺ in further analysis for this reason. While Rpd3L/ Clr6 Iʹ, Clr6 Iʺ, and Clr6S (Complex II) positively contributed to spreading (“loss” phenotype), several members of the Rpd3L-Expanded complex antagonized spreading and were found as hits inducing a “gain” phenotype (Figure 2C). This includes a subset belonging to the Set3 Complex (Set3, Hif2, Hos2, Snt1). Next, we confirmed the phenotypes of *Δfkh2* and *Δprw1* by generating de novo generated gene deletions in the MAT ΔREIII heterochromatin spreading sensor (Figure 2C). The *Δfkh2 Δprw1* double mutant displayed a similar phenotype to the single mutants, corroborating the notion that Fkh2 acts in the same pathway as Prw1 (Figure 2C).

Overall, this data suggests that Clr6 Iʹ, Clr6 Iʺ and Clr6S HDAC complexes specifically promote heterochromatin spreading in addition to their described roles in transcriptional gene silencing.

### Clr6 complexes, in contrast to Clr3, promote H3K9me2 spreading and do not contribute significantly to nucleation

Our above analysis suggested that Clr6 complex subunits contribute to the spreading of gene silencing. In what comes below, we address the mechanisms underlying this phenotype, and contrast the role of Clr6 to that of another prominent heterochromatic HDAC, Clr3. First, we examined whether the phenotype observed for Δprw1 and *Δfkh2* also holds true for clr6 itself, which codes for the catalytic subunit. To do this, we explored the *clr6-1* allele, which displays a hypomorphic phenotype at permissive temperatures (note that clr6 is an essential gene) (Grewal et al. 1998). Under this condition, *clr6-1* shows moderate spreading defects in MAT ΔREIII without affecting nucleation, which is consistent with its role in nucleation-distal silencing (**Figure 3 B vs. A**). In contrast, mutants in the HDAC Clr3 behaved differently, showing a complete loss of silencing when employing a catalytic dead version of Clr3 (*clr3-D232N*) (**Figure 3 C vs A, Figure 1**). This result implies that the two major HDACs, Clr6 and Clr3, have different roles in heterochromatin silencing.

**Figure 3:**
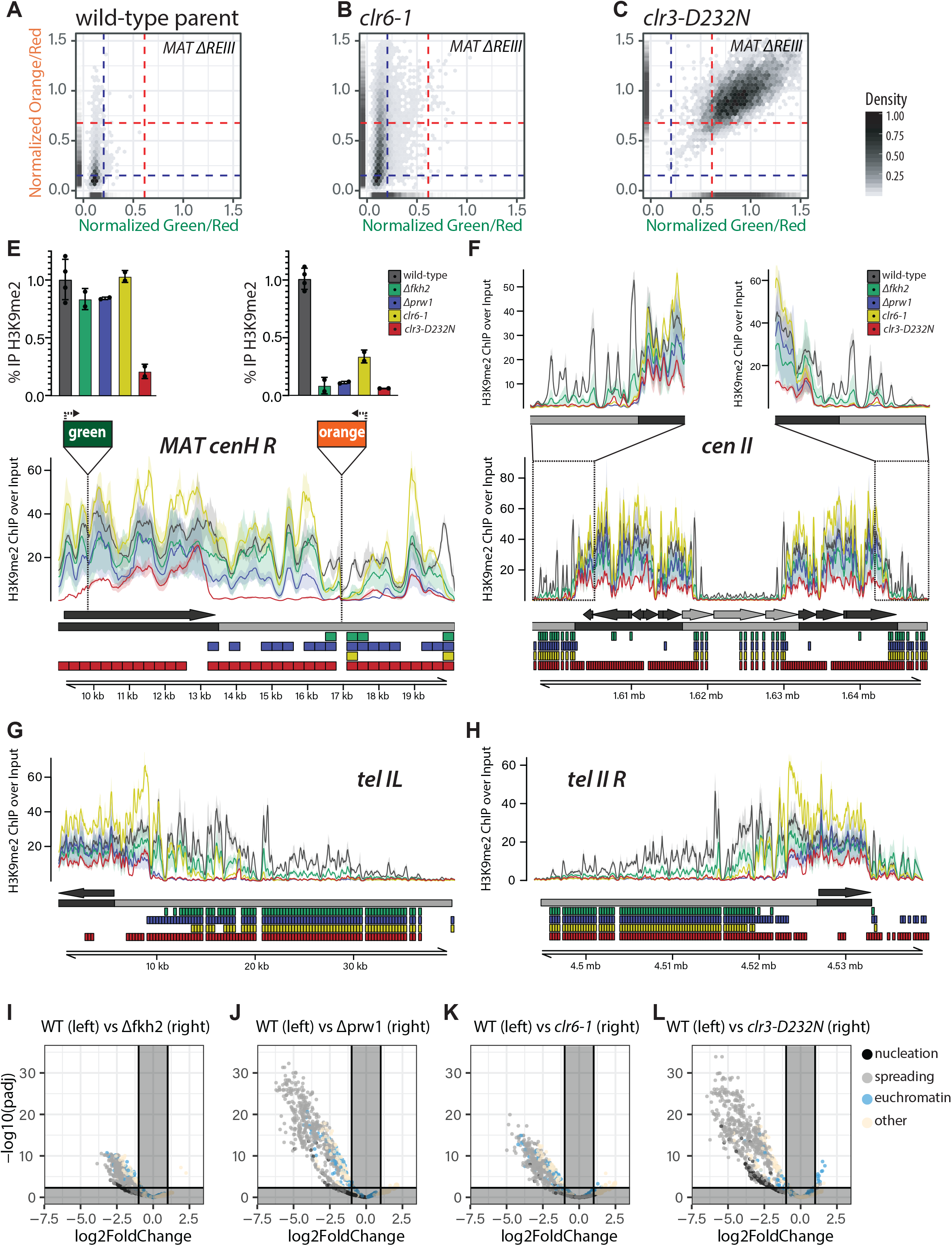
Fkh2-containing Clr6 Complexes regulate heterochromatin spreading at constitutive and facultative heterochromatin loci. **A.-C.** The hypomorphic *clr6-1* allele exhibits a loss of spreading phenotype, while the catalytic dead *clr3-D232N* allele loses all silencing. 2D density hexbin and rug plots in the *MATΔREIII* background of the parent strain (**A.**) run together with B. &C.; *clr6-1* (**B.**); and *clr3-D232N* (**C.**). A rug plot is included on the X and Y axes indicating the 1D density for each color. Rug lines are colored with partial transparency to assist with visualization of density changes. Dashed blue lines indicate the values for repressed fluorescence state and dashed red lines indicate values for fully expressed fluorescence state. **E.**-**H.** Visualization of H3K9me2 ChIP-Seq signals in the *MAT ΔREIII* background at the right side of the MAT locus (E.), centromere II (F.), and telomere and subtelomere IL and IIR (G.&H.). ChIP/Input normalized signal for 25bp intervals is plotted as mean (line) and 95% confidence interval (shade) for each genotype (see legend). Below the signal tracks the following annotations are present in order from top to bottom: (1) features of interest (i.e. nucleators, dark grey; non-nucleator, light grey) based on coordinates and strand derived from Pombase. (2) Nucleation and spreading annotation zones (based on (1)) are represented by dark grey and light grey boxes respectively. Spreading zones are defined to be between or outside of nucleation zones. (3-6) 300bp regions determined to be significantly differentially enriched for the comparisons between *Δfkh2* and wild-type (green), *Δprw1* and wild-type (blue), *clr6-1* and wild-type (yellow), *clr3-D232N* and wild-type (red) are annotated as colored boxes respectively. In E., bars above signal tracks indicate wild-type normalized H3K9me2 ChIP-RTqPCR signals for indicated genotypes conducted independently of the ChIP-Seq experiment at “green” and “orange” reporters. Error bars represent 1SD. **I.-L.** Volcano plots representing -log10(adjusted p-value) vs log2FoldChange values for mutants (I., *Δfkh2;* J., *Δprw1;* K, *clr6-1;* L., *clr3-D232N*) over WT. P-values were corrected for multiple testing with the Benjamini-Hochberg procedure. Cutoff values for adjusted p-value < 0.005 and absolute value Log2FoldChange > 1 are annotated on the plot. Dots represent individual 300bp windows tested for differential enrichment. Dots are colored by their annotation to nucleation or spreading zones, presence within a previously identified euchromatin embedded H3K9me2 heterochromatin region (“island”, “HOOD”, or “region”), or regions outside these categories (other).

Next we examined directly whether heterochromatin assembly is impacted. To this end, we focused on the H3K9me2 mark, which signals heterochromatin formation and can accumulate without major changes in gene expression (Jih et al. 2017). This allows us to examine chromatin structure independent of gene silencing. We performed H3K9me2 ChIP-Seq analysis in wild-type and the mutants *Δfkh2*, *Δprw1*, *clr6-1*, and *clr3-D232N* and analyzed the data in two independent ways: First, we produced input-normalized signal tracks, plotting mean and 95% confidence interval per genotype calculated from multiple replicates (**Figure 3 E-H**; see methods). Second, we conducted a differential enrichment analysis that examines 300bp windows along the genome containing above-background signal for significantly different accumulation of H3K9me2 between each mutant and the wild-type (**Figure 3 E-H**; see methods).

Principal Component Analysis (PCA) on the overall phenotype revealed that mutant isolates segregated from the wild-type along PC1, with Clr6 mutants diverging additionally from wild-type along PC2 (**Figure 3 Supplement 1A**). Focusing on the signal tracks as well as the differential enrichment analysis, we make the following overall observations across the genome: The three Clr6-related mutants, *Δfkh2*, *Δprw1*, and *clr6-1*, show defects in heterochromatin spreading at all heterochromatic locations in the genome (**Figure 3E-H**). This is evidenced both by separation of the 95% confidence intervals of the mutant versus wild-type signal tracks and the differential enrichment analysis, with the effect most prominent at the pericentromeres (**Figure 3F, Figure 3 Supplement 1 C, D**) and the sub-telomeric regions (**Figure 3 G,H, Figure 3 Supplement 1E,F**). At the MAT locus, the effect is more restricted to the region centromere-distal from cenH **(Figure 3E, Figure 3 Supplement 2G**), consistent with the location of “orange” (**Figure 1 Supplement 1**). This further does not appear to impact the region near REII, which is centromere-proximal from cenH. Importantly, Clr6 mutants show little defect in H3K9me2 accumulation at nucleation centers, especially those driven by RNAi, i.e. cenH at MAT, the dg and dh repeats at the pericentromere, and homologous repeats at the subtelomeric *tlh1/2* gene (**Figure 3E-H**). This suggests that Clr6 complexes largely do not contribute to heterochromatin assembly during nucleation, but instead are essential for spreading. In contrast, Clr3 appears to act upstream, as the catalytic mutant *clr3-D232N* shows strong H3K9me2 accumulation defects at all major nucleation centers as well as the distal regions (**Figure 3E-H**). This is consistent with the view that spreading is conditioned on preceding successful heterochromatin assembly at nucleation sites. The sole exception are heterochromatin islands, where surprisingly, *clr3-D232N* shows increased H3K9me2 accumulation, possibly due to redistribution form RNAi nucleation centers elsewhere (**Figure 3 Supplement 1 G-I**). At these regions, Clr6 mutants, in contrast, show loss of H3K9me2.

We further analyzed the different impact of Clr6 and Clr3 on H3K9me2 at nucleation sites and distal regions via volcano plots of regions tested for differential enrichment in comparisons of *Δfkh2*, *Δprw1*, *clr6-1,* and *clr3-D232N* each to wild-type (**Figure 3 I-L, Figure 3 Supplement 2A-D**). While all mutants have significantly reduced enrichment relative to wild-type at distal sites subject to spreading, these plots reveal key features that separate the phenotype of the Clr6C mutants from clr3-D232N: We find, (1) that nucleation center sequences are significantly reduced in the *clr3-D232N* versus wild-type comparison, but not in Clr6 complex mutant comparisons to wild-type. This further indicates that *clr3-D232N*, but not Clr6 mutants, show significant H3K9me2 accumulation defects at nucleation centers. In addition, this analysis reveals (2) that, in comparison to wild-type, *clr3-D232N* shows significantly increased H3K9me2 enrichment at euchromatic sites, namely islands, which is absent for the Clr6-related mutants (**Figure 3L, right**). Overall, these analyses reinforce the view that Clr6 primarily impacts spreading of H3K9me2, but not its accumulation at nucleation centers.

### Fkh2 is part of several Clr6 complexes in vivo

It is not established how Fkh2 integrates into Clr6 complexes, or how it may contribute to Clr6 activity. It has previously been shown that Fkh2 coimmunoprecipitates with Clr6 (Zilio et al. 2014). We validated this result here for Clr6 and Sds3, which specifies Clr6 I complexes (Figure 4 Supplement 1A). However, this result does not imply necessarily that Fkh2 is a resident member of Clr6 complexes, and if so, of which one(s). To address this question, we performed sucrose gradient fractionations of cellular lysates, similar to prior analyses (Nicolas et al. 2007), using a Fkh2-TAP fusion and epitope-tagged Clr6-13MYC. Clr6 is distributed into a number of complexes, as shown previously, but does not appear to exist in free form (**Figure 4A and Figure 4 Supplement 1C**). Fkh2-TAP is also distributed into several Clr6 peaks: It associates strongly with a large Clr6 complex focused in fraction 10. It also associates with Clr6 in fractions 5-7 but most strongly in fraction 7. Fkh2, unlike Clr6, also migrated in an apparent free form (fraction 2), consistent with its role as a transcription factor (Buck et al. 2004; Bulmer et al. 2004). When we performed this experiment with Fkh2-13MYC and Sds3-TAP, we find Fkh2 associated with the main Sds3 fractions (peaks in fractions 9-10, **Figure 4 Supplement 1B**). Hence, we believe that this peak represents the Clr6 Iʹ (Nicolas et al. 2007) and the related Clr6 Iʺ complexes (Zilio et al. 2014), which would be consistent with the involvement of Clr6 Iʺ subunits in the regulation of spreading (**Figure 2C**). Based on prior analysis (Nicolas et al. 2007), the mid-sized complex containing Clr6-13MYC and Fkh2-TAP (**Figure 4A**, fraction 7), likely, represents complex II, which is consistent with the involvement of Cph2 and Alp13 as spreading regulators across all chromatin contexts (**Figure 1, Figure 2C**). Next, we sought to address if this comigration indicates stable Clr6 complex association. We predicted that the migration pattern of Fkh2 would change in mutants of core Clr6 complex members. When we performed sucrose gradient analysis with Fkh2-13MYC (as opposed to Fkh2-TAP above) in the *Δprw1* mutant, we found that the peaks associated with bound, but not free, Fkh2-MYC are shifted towards lower molecular weight by one fraction; this was seen for both, large and small Clr6 complexes (**Figure 4B**). We note that in this experiment, more Fkh2-MYC was detected in lower molecular weight migrating complexes. Therefore, we conclude that Fkh2 is a *bone fide* member of Clr6 complexes.

**Figure 4:**
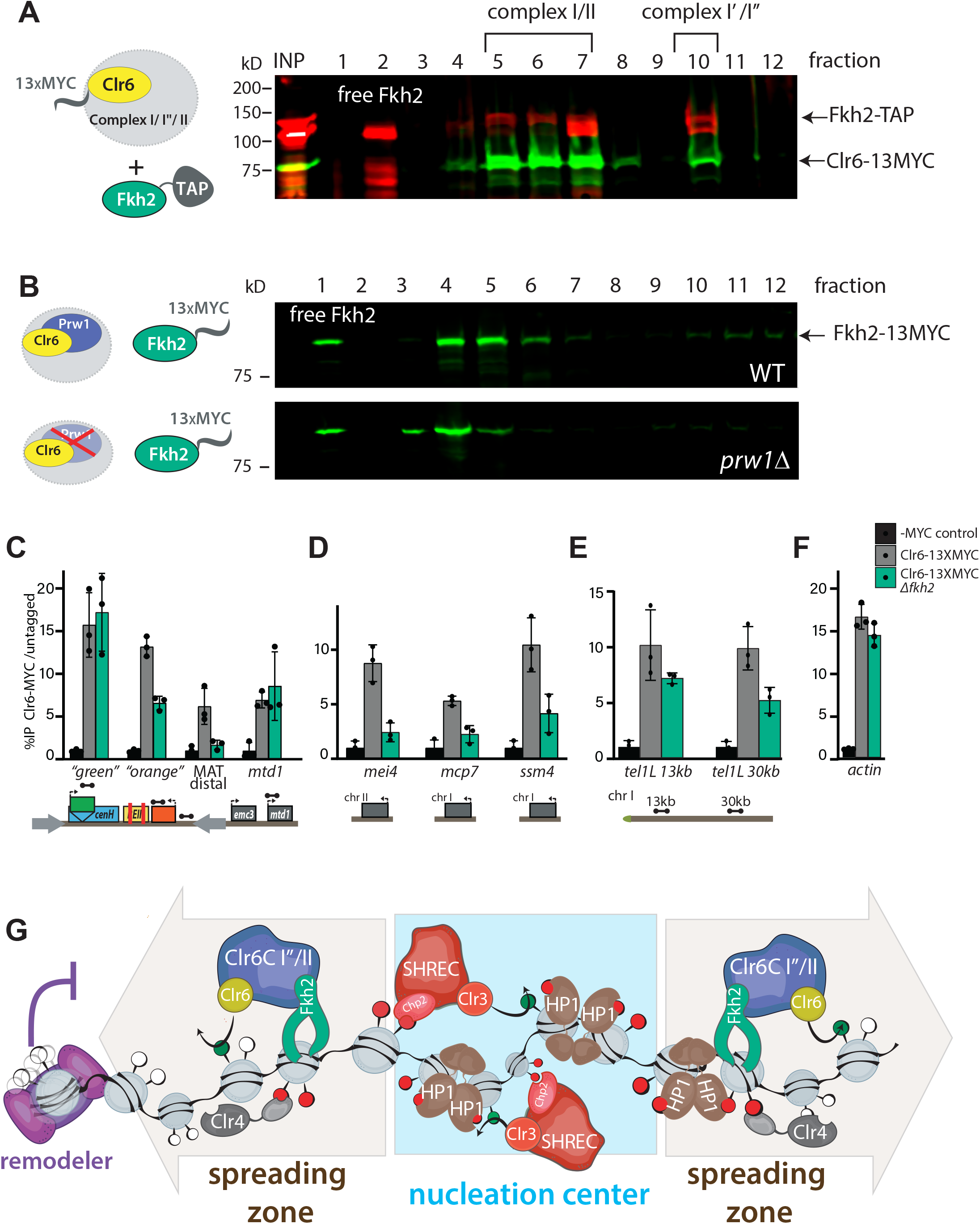
Fkh2 is a constituent member of multiple Clr6 complexes and directs Clr6 to nucleation-distal heterochromatin regions. **A.** Fkh2 co-migrates with medium sized and large Clr6 complexes. Western blot against TAP (red) and MYC (green) on fractions of a sucrose density gradient of whole cells extract containing Fkh2-TAP and Clr6-MYC. Fkh2 migrates as a free protein on top of the gradient and in both major peaks of Clr6. Complex annotation based on Nicolas et el., 2007. **B.** Fkh2 migration in sucrose gradients depends on Prw1. Western blot against MYC (green) on fractions of a sucrose density gradient of wild-type (WT) or *Δprw1* whole cell extract containing Fkh2:MYC. While the free Fkh2 fraction is not affected by the absence of Prw1, the medium and large peak fractions shift towards the top of the gradient. **C.-F.** Clr6 chromatin localization in heterochromatin spreading areas and heterochromatin islands depends partially on Fkh2. Clr6:MYC ChIP qPCR in wild-type (Clr6-13XMYC) or *Δfkh2* (Clr6-13XMYC *Δfkh2*) background relative to an untagged (-MYC) control, in *MATΔREIIII* at the HSS reporter and downstream (C.), heterochromatin islands (D.), tel1L (E.), and *act1* (F). **G.** Model for control of heterochromatin spreading by Clr3, Clr6 and chromatin remodeler complexes. The Clr3-containing SHREC complex is essential for heterochromatin assembly at noncoding RNA-driven nucleation centers, such as *cenH, dg/dh* repeats or *tlh1/2.* There, SHREC activity enables normal HP1 and Clr4 H3K9 methylase activity and is upstream to the spreading process. In the spreading zone specifically, Fkh2-containing Clr6 complexes (complex Iʺ or complex II) are required to propagate H3K9 methylation and gene silencing. Fkh2 recruits Clr6 to nucleation center-distal chromatin. Chromatin remodelers repel spreading via nucleosome destabilization, thus hinder the “guided-state” nucleosome-to-nucleosome spreading mechanism of Clr4.

### Fkh2 helps direct Clr6 to distal heterochromatin sites specifically, and not euchromatic or nucleation sites

We next sought to understand how Fkh2 helps Clr6 spread H3K9me2. One trivial possibility is that Fkh2 directs the transcription of major heterochromatin components, independent of its association with Clr6 complexes. To test this notion, we performed RT-qPCR for 9 major heterochromatin regulators, including representatives of the ClrC, RITS, Clr6, and SHREC complexes, as well as swi6. We do not observe any significant reduction of these transcripts in *Δfkh2* compared to wild-type (**Figure 4 Supplement 2**), suggesting that Fkh2 acts via another mechanism. We next tested whether Fkh2 affects the chromatin localization of Clr6. Using ChIP, we tracked the chromatin association of Clr6-13MYC at various heterochromatic loci in WT or *Δfkh2* in the MAT ΔREIII heterochromatin spreading sensor background. At the MAT locus, Clr6-13MYC was efficiently detected above untagged background signal at cenH “green” or at mtd1, a gene in euchromatin outside the IR-R MAT boundary. At these loci, Clr6 recruitment was not affected by the absence of Fkh2 (**Figure 4C**). However, at “green”-distal sites, namely the “orange” reporter and a more boundary proximal site, deletion of *fhk2* reduces the localization of Clr6. Similarly, *Δfkh2* affects Clr6 localization at the heterochromatin islands *mei4, mcp7*, and *ssm4* (**Figure 4D**). At telomeres, where *Δfkh2* also has a significant impact on H3K9me2 spreading, we also find a significant reduction in Clr6 chromatin association at distal sites (**Figure 4E**, 30kb) but not at the euchromatic control locus, act1 (**Figure 4F**). Therefore, it appears that one role of Fkh2 in promoting heterochromatin spreading is to strengthen the recruitment of Clr6 to nucleation-distal heterochromatic sites.

## Discussion

The formation of a heterochromatin domain requires three interconnected steps, nucleation, assembly of heterochromatin structures, and the lateral spreading from DNA-sequence driven nucleation sites. The genetic circuitry that regulates spreading at different chromatin loci is not well understood. Here, our ability to separate requirements for nucleation and distal spreading within heterochromatin domains allowed us to pinpoint which factors are necessary to drive or restrain spreading at different genomic loci. A key finding from this work is the requirement of variants of the Clr6 HDAC complex specifically in the spreading reaction, in addition to the antagonism by a broad class of chromatin remodelers. Additionally, we find that different chromatin contexts have specific requirements for spreading, for example, the role of SAGA in promoting spreading within euchromatin.

Of the HDACs involved in heterochromatin function, Clr3 is likely the best described. Its associated SHREC complex is required for silencing, likely via its ability to repress nucleosome turnover (Aygun et al. 2013), maintain nucleosome occupancy (Sugiyama et al. 2007; Garcia et al. 2010), and remove H3K14 acetylation known to antagonize heterochromatin assembly (Wirén et al. 2005). Our results show that Clr3 catalytic and deletion mutants completely lose heterochromatic silencing and are defective in H3K9me2 accumulation (**Figure 1, 3**) including at nucleation centers (**Figure 3, Figure 3 Supplement 3**). Similarly, the Sir2 HDAC is broadly implicated in heterochromatin nucleation and assembly (Shankaranarayana et al. 2003; Alper et al. 2013), and we also observe near-complete silencing loss in *Δsir2* (**Figure 1 Supplement 5**). In contrast to Clr3 and Sir2, Clr6 is required specifically for heterochromatin spreading, but not assembly at nucleation centers (**Figures 1-3**). This is consistent with the finding that the clr6-1 allele has only small impacts on transcription of the cenH nucleator-encoded ncRNAs (Yamane et al. 2011). Our screen showed that several, but not all members, of the recently described Clr6 complex Iʺ (Zilio et al. 2014) promote spreading, Key subunits of Clr6 II and Clr6 Iʹ were also featured as hits. That not all annotated Clr6 subunits share gain or loss of spreading phenotypes may suggest these subunits do not contribute to heterochromatin spreading, but instead mediate other functions of the complex. On the other hand, Fkh2 is a very prominent subunit promoting spreading, especially at MAT. As we find that Fkh2 is a resident member of several Clr6 complexes, the Iʹ,Iʺ, and II types (**Figure 4A, B**), we believe these results indicate two possible, nonexclusive interpretations: (1) the composition of different Clr6 subcomplexes in vivo is either more dynamic than previously thought, and/or (2) a number of different, preassembled Clr6 complexes can associate with Fkh2, which imparts a role in spreading regulation. Interestingly, the Set3-submodule that typifies the Rpd3L-Expanded complex (Shevchenko et al. 2008) has a distinct spreading-antagonizing behavior (**Figure 2C**). This contrasts with a mild positive role of the Set3 complex at pericentromeres (Yu et al. 2016). Taken together, we believe our data suggest that Clr3 works upstream, to establish heterochromatin at noncoding-RNA driven nucleation centers, and Fkh2-containing Clr6 complexes function downstream to spread heterochromatin structures outward (**Figure 4G**).

What mediates the spreading-specific role of Clr6 complexes? One possible explanation is that they act in conjunction with the histone chaperone Asf/HIRA which cooperates with Clr6 in gene silencing at ncRNA nucleators (Yamane et al. 2011). However, we do not favor that this pathway mediates distal spreading, since Asf/HIRA subunits Hip1, Hip3 and Slm9 have mild or no phenotype for spreading in MAT contexts. Asf/HIRA mutant phenotypes were more pronounced in ECT, a context that is less reliant on Clr6 for spreading (**Figure 1**, see below). We believe Fkh2 plays a key role in imparting this spreading-specific role for Clr6. Fkh2 is a transcription factor that regulates meiotic genes in *S. pombe* (Buck et al. 2004; Bulmer et al. 2004; Alves-Rodrigues et al. 2016). However, the observation that loss of Fkh2 impacts H3K9me2 spreading at all sites (**Figure 3, Figure 3 Supplement 3**) makes it less likely in our view that Fkh2 acts via binding to canonical sequence motifs. Fkh2 has also been shown to act as a chromatin organizing protein through clustering origins of replication (Knott et al. 2012). It is conceivable that Fkh2 creates chromatin environments that are conducive to nucleation -distal Clr6 recruitment, either by tethering to a nuclear compartment such as the nuclear periphery (Holla et al. 2020) or via chromatin conformational or biophysical changes (Sanulli and G 2020; Zenk et al. 2021). In either case, Fkh2 promotes the recruitment of Clr6 complexes to the spreading zone (**Figure 4C-E).** The spreading-specific role of Clr6 complexes may be additionally supported by other known recruitment mechanisms, for instance HP1/Swi6 (Hall et al. 2002; Canzio et al. 2011) (Fischer et al. 2009). While the precise mechanism of Fkh2-mediated Clr6 recruitment will be the subject of further study, our results unambiguously demonstrate that Fkh2 typifies a specific functional mode of the conserved HDAC Clr6/RPD3 among its numerous essential activities in gene regulation (Yang and Seto 2008).

Our results show a role of chromatin remodelers across several classes in antagonizing spreading. This role is widespread, as Ino80, Swr1C, SWI/SNF and RSC appear to contribute to antagonizing heterochromatin spreading. This broad antagonism contrasts with more specific functions uncovered previously for Ino80/Swir1C (Meneghini et al. 2003). As above for Clr6, not all subunits show a spreading phenotype. Remodelers have been implicated in negatively regulating heterochromatin function by creating specific nucleosome free regions (NFRs, (Lorch and Kornberg 2017)) that antagonize heterochromatin. Since NFRs may be roadblocks to spreading (Garcia et al. 2010; Lantermann et al. 2010), it is possible that remodelers employ this mechanism to restrain heterochromatin spreading globally. In addition, remodelers such as SWI/SNF and RSC destabilize nucleosomes generally (Narlikar et al. 2001; Rowe and Narlikar 2010), leading to increased turnover (Rawal et al. 2018), which would antagonize heterochromatin spreading. This increased turnover may be tolerated at ncRNA nucleation sites, where turnover is at near euchromatic levels (Greenstein et al. 2018), likely due to ncRNA transcription (Volpe et al. 2002; Noma et al. 2004). This would suggest that regulation of nucleosome stability has a particular significance at distal, but not nucleation sites.

Beyond the broad finding of Clr6 and remodelers as specific promoters and antagonizes of spreading respectively, this study uncovered several locus- and nucleation center type-specific pathways. Here we would like to highlight two main observations:

1. Distinct factors are required for similar nucleators in different chromatin environments. ECT and MAT ΔREIII are both driven by related ncRNA nucleators (dh and cenH, respectively) and have remarkably similar behaviors with respect to nucleation and spreading across the cell population (Greenstein et al. 2018). Efficient spreading specifically at ECT requires Hip1, and moderately Slm9, which code for a key subunits of the HIRA H3/H4 chaperone. HIRA has been implicated in stabilizing heterochromatic nucleosomes (Yamane et al. 2011). Hence, given that transcribed chromatin is known to destabilize nucleosomes, it seems likely that this specific requirement reflects the challenge faced by heterochromatic domains when expanding within gene-rich chromatin. ECT is also particularly reliant on the SAGA complex for spreading **(Figure 2A, Figure 2 Supplement 2**). This initially may seem counterintuitive, as SAGA has been shown to be recruited by Epe1 to antagonize heterochromatin assembly at constitutive sites (Bao et al. 2019). In fact, in our screen SAGA plays a less prominent role in the MAT context. This requirement for SAGA at ECT may be connected to the observation that SAGA can modulate the chromatin recruitment of remodelers, such as SWI/SNF, via direct acetylation (Kim et al. 2010). Therefore, one possible explanation that remains to be tested for the SAGA phenotype we observe is acetylation of SWI/SNF and possibly other remodelers that antagonize spreading, releasing them from chromatin.
2. Spreading from qualitatively different nucleators within the same environment, namely REIII and cenH, also differ in their sensitivity to different mutants. The significant overlap in factors between WT MAT and MAT ΔREIII indicates that heterochromatin formation at MAT is dominated by the ncRNA nucleator cenH, in agreement with our previous findings (Greenstein et al. 2018). The REIII element, which nucleates heterochromatin independent of ncRNA (Jia et al. 2004), has different requirements. For example, ncRNA-independent spreading at REIII (MAT ΔcenH) is uniquely promoted by the MTOR pathway Gad8 kinase, partially consistent with a previous report implicating Gad8 for MAT silencing (Cohen et al. 2018). While Gad8 is reported to target Fkh2 for phosphorylation, *Δfkh2* has very weak effects on MAT ΔcenH. Other potential phosphorylation targets of Gad8, if it acts through its kinase function, in promoting spreading from REIII remain to be established. We note that REIII can confer a high propensity for local intergenerational inheritance of silencing (Grewal and Klar 1996; Greenstein et al. 2018). Therefore, a formal possibility for spreading defects in the MAT ΔcenH context, or others with high intergenerational stability, is that the gene identified is required for the maintenance of heterochromatin outside the nucleator, rather than its initial formation by spreading.

In this work, we defined how regulation of heterochromatin silencing and nucleation differ in fundamental ways from distal spreading. While similar nucleation elements likely rely on a common set of machinery, the success of heterochromatin spreading appears much more sensitive to the chromatin context, particularly in gene-rich versus gene-poor chromatin. Our findings will likely have important implications for directing gene silencing during cellular differentiation. In this situation, regions that have previously been in a transcriptionally active state are invaded by heterochromatin and will have to compete for core spreading factors in a dosage limited system (Eissenberg et al. 1992; Nakayama et al. 2000). We note that several of the factors we identify as critical to regulating spreading in euchromatinic environments are conserved in metazoans, opening the possibility that they contribute differentiation via heterochromatin control in these organisms.

## Methods

### Mutant Generation for Genetic Screen

We generated a 408 gene deletion mini-library that represent a subset of the S. pombe Bioneer deletion library, focused on nuclear function genes. The library (table 1) was assembled via three criteria: 1. ∼200 genes coding for proteins that evidence “nuclear dot” morphology based on a high-throughput YFP-tagged“ORFeome” screen (Matsuyama et al., 2006, also used in Braun et al., 2011). 2. ∼50 deletions of central *S. pombe* DNA binding transcription factors. 3. ∼150 gene deletions selected based on a search of annotations for chromatin regulation-related functions (majority of this set), or prior preliminary data for roles in regulation of heterochromatin function. Deletions that grow poorly in rich media were eliminated. Several Bioneer collection mutants were independently validated and produced de novo. For the ectopic locus HSS reporter strain, the screen was performed essentially as described (Greenstein et al. 2019). Briefly, the parent HSS reporter strain was crossed to the library. Crosses were performed as described (Verrier et al. 2015; Barrales et al. 2016; Greenstein et al. 2018; Greenstein et al. 2019) using a RoToR HDA colony pinning robot (Singer). For the MAT HSS reporter strains, the screen was performed essentially as described (Greenstein et al. 2019) with the exception that crosses were generated using a 96 well manual pinner. Note that the MAT ΔcenH, we used an OFF isolate as described (Greenstein et al. 2018). This is because ΔcenH behaves in a bimodal fashion, producing stable ON (no heterochromatin) and OFF (heterochromatin) alleles at MAT. In addition, three *Δclr4* mutant isolates and six individual parent isolates from each genomic context were included as controls. Crosses for the ectopic HSS strains were performed using SPAS media for 4d at room temperature, while for the MAT HSS strain crosses were performed on ME media for 3d at 27°C. For all strains, crosses were incubated for 5d at 42°C to retain spores, while removing unmated haploid and diploid cells. The ectopic HSS spores were germinated on YES medium supplemented with G418, hygromycin B, and nourseothricin. For MAT HSS strains, spores were germinated on YES medium supplemented with G418 and hygromycin B. The resulting colonies were pinned into YES liquid medium for overnight growth and then prepared for flow cytometry as described below.

### Flow Cytometry Data Collection and Normalization for Genetic Screen

In preparation for flow cytometry, overnight cultures were diluted to OD = 0.1 (approximately a 1:40 dilution) in rich media (YES) and incubated at 32 C with shaking of rpm for 4–6 hours. For the ectopic locus HSS strains, flow cytometry was performed essentially as described (Greenstein et al. 2018; Greenstein et al. 2019). For the MAT locus HSS strains, flow cytometry was performed using a Fortessa X20 Dual instrument (Becton Dickinson) attached with high throughput sampler (HTS) module. With a threshold of 30,000 events, samples sizes ranged from ∼1000 to 30,000 cells depending on strain growth. Fluorescence detection, compensation, and data analysis were as described. The R scripts for analysis is included as a text file. Flow data deriving from the genetic screen for individual strains are represented as 2d density hexbin plots in Figure 1, 1S3, 1S5, 2S1, and 2S3. Dashed red and blue guidelines respectively indicate median minus 2SD of “green”/“orange”-On cells (*Δclr4* control cells with normalized fluorescence above 0.5) and median plus 2SD of “green”/ “orange”-Off cells (“red”-Only control).

We note that an error occurred relative to v1 of this manuscript: .fcs files in the 3.1 version are exported with a color compensation matrix appended to each sample but not applied, in contrast to .fcs 2.0 files where compensation is applied upon export. Thus, since our R scripts assumed color compensated data to be exported, no additional compensation was applied. We noted this error during revisions and have updated our R scripts to account for this and process color compensation. We then applied a uniform 2 standard deviation threshold for all of our controls and normalization. Note that the numerical value for some spreading hits changed as a result.

### Spreading Analysis

Nucleated cells were extracted using a “green”-off gate, using median of a “red”-only control plus 2 times the SD. Enrichment of cell populations in particular “orange” fluorescence ranges (Gridn) are calculated as Grid ^mut/par^: fraction of mutant population is divided by the fraction of parent population in one grid. The intervals of “orange” fluorescence used in grids are determined by: median plus 2SD of “orange”-Off cells (“red”-Only control), median minus 2 SD of “orange”-On cells (Δclr4 control cells with normalized fluorescence above 0.5) and the median of the two. For ECT, instead of using a “red”-Only control strain, we adjusted the fluorescence value of a fluorescence-negative (unstained) strain by the “red” fluorescence from the *Δclr4* control. To evaluate gain of spreading phenotype, enrichment in Grid 1 in WT MAT, MAT ΔREIII and ECT were calculated. To evaluate loss of spreading phenotype, enrichment in Grid 3 and 4 in MAT ΔcenH and WT MAT as well as Grid 4 in MAT ΔREIII and ECT were calculated. The distribution of the Gri_dn_^mut/par^ were plotted as swarmplots with annotation of 85th percentile and median plus 2SD of parent isolates Gridnmut/par. Gene hit lists comprised mutations above median and 2SD within the 85th percentile. Upset plots were generated using the R package UpSetR (Conway et al. 2017). Beewarm plots were plotted using the R packages ggbeeswarm (Clarke 2017)

### GO Complex and Sub-Complex analysis

#### generating the heatmap count data

GO Complexes – Based on the GO Complex annotations [link] retrieved from pombase (Lock et al. 2018; Consortium 2019), GO complex membership was determined for genes identified as hits for each strain background and hit category (gain/loss). Briefly, using functions from the R package dplyr (Wickham H., François R., Henry L. and Müller K. (2020). dplyr: A Grammar of Data Manipulation. R package version 0.8.4. https://CRAN.R-project.org/package=dplyr), gene names were converted to systematic ID numbers and these systematic IDs were queried against the GO complex annotation table. The number of times a GO complex appeared per background and hit category was tabulated. Genes can be associated with any number of GO complexes depending on their annotations. However, any particular gene was only counted once per GO complex despite potentially being annotated to that GO complex by more than one evidence code. The unique list of GO complexes for all hits was determined and a matrix was computed representing the number of times each GO complex (row) was identified per strain/hit category (column). This counts matrix was used to generate the GO complex heatmap in Figure 2S1, described below.

Hit tables – Genes annotated to the seven complexes in Figure 2A,B,C were obtained from pombase (Lock et al. 2018). fkh2 was added to the Clr6 Iʺ complex given the protein contacts described previously (Zilio et al. 2014). For the unique set of genes per panel it was determined if each gene was identified as a hit in each strain background/hit category combination. The data was summarized in a counts matrix where rows represent the unique list of genes per panel and columns represent the strain background / hit category. The counts matrix for each set of genes was used to generate the heatmaps in Figure 2A,B, C as described below.

#### generating the heatmap clustering

Using the R package ComplexHeatmap (Gu et al. 2016), both row and column dendrogram and clustering were generated using hierarchical clustering. Based on an optimal Silhouette score, the strain background / hit category (columns) were clustered into 2 clusters. The dendrogram representing complexes (Figure 2S1) or genes (Figure 2A,B, C) in rows were not separated because validations of the clustering by connectivity, Dunn index or Silhouette score were inconclusive. Clustering validations were conducted using the R package clValid (Brock, G., Pihur, V., Datta, S. and Datta, S. (2008) clValid: An R Package for Cluster Validation Journal of Statistical Software 25(4) URL: http://www.jstatsoft.org/v25/i04).

### Validation Strain and Plasmid Construction

Plasmid constructs for gene knockout validation were generated by in vivo recombination as described (Greenstein et al. 2018; Greenstein et al. 2019). S. pombe transformants were selected as described (Greenstein et al. 2018). For microscopy, hygMX super-folder GFP (SFGFP) constructs for C-terminal tagging we described previously (Al-Sady 2016) were amplified with 175bp ultramer primers with homology to apm3 or apl5 and transformed into a Swi6:E2C kanMX strain. Apm3:SFGFP;Swi6:E2C and Apl5:SFGFP;Swi6:E2C strains were selected on hygromycin B and G418. Integrations and gene knockout were confirmed by PCR.

### Flow Cytometry Data Collection and Normalization for Validation

For validation flow cytometry experiments, cells were grown as described (Greenstein et al. 2018; Greenstein et al. 2019) with the exception that cells were diluted into YES medium and grown 5-8 hours before measurement. Flow cytometry was performed as above. Depending on strain growth and the volume collected per experiment, fluorescence values were measured for ∼20,000-100,000 cells per replicate. Fluorescence detection, compensation, and data analysis were as described (Al-Sady et al. 2016; Greenstein et al. 2018; Greenstein et al. 2019) with the exception that the guide-lines for boundary values of “off” and “on” states were determined using median of a Red-Only control plus 3 times the median absolute deviation (MAD) and median of *Δclr4* minus 2 times the MAD value respectively. Validation flow cytometry plots were generated using the ggplot2 R package (Wickham 2016).

### Chromatin Immunoprecipitation and Quantification

Chromatin Immunoprecipitation (ChIP) was performed essentially as described (Greenstein et al. 2018; Greenstein et al. 2019) Bulk populations of cells for were grown overnight to saturation in YES medium. For anti-H3K9me2 ChIP, the following morning, cultures were diluted to OD 0.1 in 25mL YES and grown for 8h at 32°C and 225rpm. Based on OD measurements, 60×106 cells were fixed and processed for ChIP as previously described (Greenstein et al. 2018) without the addition of W303 carrier. For anti-MYC ChIP, 40mL cultures were grown to OD 0.4-0.7 and then incubated 30min at 18°C. Based on OD measurements, 100×106 cells were crosslinked with 1% formaldehyde at 18°C. Anti-MYC cells were lysed as described (Greenstein, 2018), except that cells were bead mill homogenized for 9 cycles. Cleared chromatin for both anti-H3K9me2 or anti-MYC ChIP samples was incubated with either 1μL of anti-H3K9me2 antibody (Abcam, ab1220) or 2μL anti-MYC antibody (Invitrogen, MA1-980, lot VL317116) overnight after a small fraction was retained as Input/WCE. DNAs were quantified by qPCR. For H3K9me2, percent immunoprecipitation (%IP, ChIP DNA/Input DNA) was calculated as described (Greenstein et al. 2018), except for Figure 2 Supplement 2, where a ratio of %IP queried locus/%IP act1 is plotted. For anti-MYC ChIP, enrichment is presented as the ratio of %IP in PAS867 or 868 (Clr6:13XMYC in PAS 332 WT or °fkh2) over %IP in PAS332 (untagged). Data was plotted in Prism (GraphPad). For comparison of different preparations of ChIP samples, %IP of mutant divided by %IP of wildtype was calculated.

### ChIP-Seq Data Collection, library preparation and sequencing

ChIP was performed essentially as above, with the following exceptions: From 60 mL cultures, 300×106 cells in logarithmic phase were fixed and processed. Sheared chromatin samples were not pre-cleared with Protein A Dynabeads, and the chromatin was directly treated with 2μL of anti-H3K9me2 antibody (Abcam 1220, Lot GR3308902-4). Barcode-indexed sequencing libraries were generated from reverse-crosslinked ChIP-DNA samples using a Kapa Hyper DNA Library Preparation Kit (Kapa Biosystems-Roche, Basel, Switzerland) and NextFlex UDI adapters (PerkinElmer, Waltham, MA). The libraries were amplified with 16 PCR cycles and cleaned with SPRI bead protocol according to the instructions of the manufactures. The fragment lengths of the sequencing libraries were verified via micro-capillary gel electrophoresis on a LabChip GX Touch system (PerkinElmer). The libraries were quantified by fluorometry on a Qubit instrument (LifeTechnologies, Carlsbad, CA), and combined in a pool at equimolar ratios. The library pool was size-selected for library molecules in the lengths of 200 to 450 bp using a Pippin-HT instrument (Sage Science, Beverly, Massachusetts). The success of the size-selection was verified on a Bioanalyzer 2100 instrument (Agilent, Santa Clara, CA). The pool was quantified with a Kapa Library Quant kit (Kapa Biosystems-Roche) on a QuantStudio 5 real-time PCR system (Applied Biosystems, Foster City, CA) and sequenced on a Illumina NextSeq 500 (Illumina, San Diego, CA) run with paired-end 40 bp reads.

### ChIP-Seq Data Analysis

Data processing for ChIP-seq analysis was performed as follows. Trimming of sequencing adaptors and sliding window quality filtering were performed using Trimmomatic v0.39 (Bolger et al. 2014). Filtered and trimmed paired-end (PE) reads were aligned to the S. pombe genome (Wood et al. 2002) with Bowtie2 v2.4.2 (Langmead and Salzberg 2012) using standard end-to-end sensitive alignment. An additional 6bp was trimmed from the 5’ end of each read prior to alignment. Sorted, indexed bam files were generated using SAMtools v1.12 (Li et al. 2009). Duplicate reads were marked with Picard tools v2.25.2 “MarkDuplicates” command. Filtered bam files were generated with SAMtools “view” with the following flags [-bh -F 3844 -f 3 -@ 4 I II III mating_type_region] to retain only properly paired reads on the listed chromosomes/contigs, and remove duplicate reads. The resulting filtered bam files were sorted and indexed with SAMtools and used for downstream genome-wide analysis. Input normalized BigWig files for signal tracks for 25bp bins were generated from the filtered bam files with the bamCompare function from deeptools v3.5.1 (Ramírez et al. 2016) using the following flags [--outFileFormat bigwig --scaleFactorsMethod readCount --operation ratio --pseudocount 1 --extendReads --samFlagInclude 64 --skipZeroOverZero --binSize 25 --numberOfProcessors 4 --effectiveGenomeSize 12591546 –exactScaling]. Fragments were counted once by including only the first mate of each pair and extending to the fragment size. For each genotype, 2 or 3 biological replicates were processed for downstream analysis. Initial whole genome clustering analyses on ChIP and Input samples and inspection for low signal to noise ratio in IGV prompted us to remove two outlier samples, one clr6-1 and wild-type respectively.

BigWig files were imported into R v4.0.3 with rtracklayer v1.50.0 (Lawrence et al. 2009). BigWig files were used to generate signal tracks comprised of the mean and confidence interval for each genotype in custom genome browser plots generated with the DataTrack() command from the Gviz Bioconductor package v1.34.1 (Hahne and Ivanek 2016). As described previously (Greenstein et al. 2019), gene annotations were imported from PomBase (Lock et al. 2018)and converted to genomic coordinates in R v3.5.1 with the makeTxDbFromGFF function from GenomicFeatures v1.32.3 (Lawrence et al. 2013) and saved out to an sqlite file. This sqlite file was imported into R v4.0.3 and used to generate feature annotations for signal tracks in Gviz with the AnnotationTrack() command.

Reads from the filtered bam files for ChIP samples were counted into either 300bp windows or 5kb bins using windowCounts() function from the Bioconductor package csaw v1.24.3 (Lun and Smyth 2016). Regions +/-1.5kb surrounding the following features (*ade6, ura4, fkh2, prw1, mat1-Mc, mat1-Mi*) were blacklisted as they represent experimental artifacts – ade6 and ura4 because their promoter and terminator regions are present in the reporter cassettes, fkh2 and prw1 because these gene’s ORFs were entirely removed in certain genetic backgrounds and mat1 genes because these regions are expressed but homologous to those found in the heterochromatic MAT locus. Global background was determined from 5kb bin count matrices and interpreted onto the 300bp windows with the csaw filterWindowsGlobal() function. Only 300bp windows where the abundance exceeded a filtering threshold of 1.7 times the global median were retained resulting in 1500 windows for further analysis.

Differential Enrichment analysis was performed on these 1500 300bp windows using the DESeq2 Bioconductor package v1.30.1 (Love et al. 2014). Size Factors were calculated on the count matrices in the global 5kb bins with the estimateSizeFactors() function and applied on the global enrichment filtered 300bp windows. The model matrix for the experimental design was constructed based on genotype. Normalized counts per bin were obtained with the vst() function from DESeq2 with the parameter blind = FALSE. VST transformed counts were used as an input to principal component analysis via the prcomp() R function on the top 500 most variable bins. PCA plots were generated with ggplot2 v3.3.3 (Wickham 2016). Differential Enrichment analysis, including estimating dispersions and fitting of a negative binomial generalized linear model, was performed with the DESeq() function. Results for pairwise contrasts between genotypes were extracted as GRanges objects with the results() function. Each of the 1500 windows was annotated as belonging to global or heterochromatin location specific nucleation/spreading/euchromatic/other categories. Volcano plots were generated with ggplot2 for each comparison by plotting -log10(padj) against log2FoldChange values. Each dot represents a 300bp region tested for differential enrichment. Dots are colored by their location annotation. Coordinates for nucleator regions are derived from feature coordinates from PomBase adjusted to a multiple of 300 so that a 300bp bin can only be annotated to one category of feature. Coordinates for spreading are defined to be between or outside of nucleator regions. Euchromatic regions are defined as coordinate ranges identified as an “island” (Zofall et al. 2012), “HOOD” (Yamanaka et al. 2013), or “region” (Wang et al. 2015)as delineated in Supplemental Table S6 from (Parsa et al. 2018). A threshold for significance of padj (Benjamini-Hochberg adjusted p-value) < 0.005 and abs(log2FoldChange) > log2(2) was applied on the results of each comparison and these cutoff values are additionally annotated on the volcano plots. For each comparison, the number of significant regions of each category was tabulated. Regions called as significant for the comparison of each mutant to WT are annotated on the custom genome browser plots generated with the Gviz AnnotationTrack() command. A Venn diagram of regions where WT signal significantly exceeds signal from mutants was generated outside of the conda environment in R v4.0.5 with the venn R package v1.10.

An additional custom analysis for the MAT locus was performed starting at the genome alignment stage. A fasta file that includes the inserted “green” and “orange” color cassettes and intact atf1/pcr1 binding sites was used was used to build a genome index for bowtie2. Alignment to this custom reference was performed with bowtie2 as above.

Alignments were filtered with SAMtools to retain only properly paired reads. Multimapping and low-quality alignments were removed with a mapq filter of -q 10. In the context of this custom reference sequence, multimapping reads represent regions that align to ura4p, ade6p, and ura4t which are present at XFP reporter cassettes, IR-R and IR-L repetitive regions, and parts of the mating type cassettes. Duplicate reads were marked and removed with Picard tools. Sorted, and indexed bam files were generated with SAMtools. BigWig files for coverage signal tracks at 10bp resolution were generated from ChIP files using the deeptools bamCoverage function with the following flags [--outFileFormat bigwig --scaleFactor ## --extendReads --samFlagInclude 64 --binSize 10 --numberOfProcessors 4 --blackListFileName cenH_blacklist.bed --effectiveGenomeSize 19996 --exactScaling]. Bam files were scaled by a custom scaling factor (replacing ## above) that adjusts for the read count in the bam file from each sample’s the full genome alignment to the read count of the bam file with the smallest number of reads in the full genome alignment. The regions included in the cenH element were blacklisted in this step as they are homologous to many sequences found at the centromeres and telomeres and in this analysis represent aggregate signal from all these regions. The resulting coverage BigWig files were used to generate signal tracks in R/Gviz as described above.

### Microscopy

Swi6:E2C; Apl5:SFGFP and Swi6:E2C; Apm3:SFGFP cells were grown is YES media as described. Slides (ibidi, Cat. No. 80606) were precoated with 100 mg/mL lectin (Sigma-Aldrich, Cat. No. L1395) diluted in water by adding lectin solution to slide for 1 min. and removing supernatant. Log-phase growing cells were applied to the slide and excess cells were rinsed off with YES. Cells were immediately imaged with a 60x objective (CFI Plan Apochromat VC 60XC WI) on a Nikon TI-E equipped with a spinning-disk confocal head (CSU10, Yokogawa) and an EM-CCD camera (Hammamatsu). Cells were imaged in brightfield and additionally excited with 488nm (SFGFP) and 561nm (E2C) lasers. Emission was collected using a 510/50 band-pass filter for GFP emission and a 600/50 band-pass filter for E2C emission. For the SFGFP and E2C channels, z-stacks were obtained at 0.3µm/slice for 11 slices total. An overlay of the maximum z-projections for SFGFP and E2C channels are shown separately from the brightfield images. Brightness and contrast were adjusted in ImageJ to clearly show both Swi6 and Apl5/Apm5 signals in the overlay. At least two isolates were imaged to confirm localization patterns.

### Reverse transcription qPCR validation of context-specific spreading mutants

For validation of context-specific spreading hits, saf5, eaf6, pht1, hip1 and gad8 mutants were crossed to PAS217 (WT-MAT), PAS332 (MAT-EIII), PAS482 (MAT-cenH), and PAS231 (ECT), respectively and mating products selected for KAN and HYG (PAS217, PAS332, and PAS482) or KAN, HYG, and NAT resistance (PAS231). For analyzing transcriptional regulation of heterochromatin regulators via Fkh2, PAS332 or PAS798 (fkh2 in PAS332) were grown as above. Two independent isolates of each strain were grown in 200 l YES in 96 well plates to log phase (OD∼0.4-0.8), washed with water and flash frozen. Total RNA was extracted from cell pellets as described (Greenstein et al 2018) using the Masterpure yeast RNA extraction kit (Lucigen). cDNA was produced from 2-3 g total RNA as described (Greenstein et al 2018) using a dT primer and Superscript IV (Invitrogen) reverse transcriptase, followed by an RNaseH step to remove RNA:DNA hybrids. qPCR was performed with primers against act1, SF-GFP and mKO2, or amplicons for heterochromatin regulator transcripts indicated in Figure 4 Supplement 2. For act1 qPCR, the cDNA was diluted 1:60. qPCR was performed as described (Greenstein et al 2019).

### Sucrose gradient analysis

Sucrose gradient analysis was performed essentially as in (Nakayama et al. 2003), with several modifications. 50mls of PAS833 (Fkh2:13XMYC), PAS836 (Fkh2:TAP; Clr6:13XMYC), or PAS837 (Fkh2:13XMYC; Sds3:TAP) were grown in YES to OD∼ 1, spun and washed with STOP buffer (150mM NaCl, 50mM NaF, 10mM EDTA and 1mM NaN3) and flash frozen. Cells were resuspended in 3ml ice cold HB-300 (50mM MOPS pH7.2, 300mM NaCl, 15mM MgCl2, 15mM EGTA, 60mM glycerophosphate, 0.1mM Na3VO4, 2mM DTT, 1% Triton X-100, 1mM PMSF, 200μM phenantroline, pepstatin A, leupeptin, aprotinin and 1X of EDTA-free protease-inhibitor cocktail (Roche)) and then lysed in the presence of 600μl 0.5mm zirconia beads (RPI) for 6X 1min, with 5 min rest on ice in between cycles, at the maximum setting in a bead mill homogenizer (Beadruptor-12, Omni International). The lysate was clarified by spinning at 18,000 x g for 20min. 500μl clarified lysate was applied to a 4-20% sucrose gradient in gradient buffer (50mM Tris-HCl pH 7.5, 50mM KCl, 1mM EDTA, 1mM DTT, 1mM PMSF, 200μM phenantroline, pepstatin A, leupeptin, aprotinin and 1X of EDTA-free protease-inhibitor cocktail (Roche)) and spung for 20hrs at 151,000 x g (r_av_) at 4°C in a swinging bucket rotor (Beckman SW-41 Ti). 12 1ml fractions were taken off the gradient and incubated for 10min at RT with 100μl 0.15% deoxycholine. Proteins were then precipitated by addition of 100 l of 50% Trichloroacetic acid and incubation on ice for 30min. Precipitates were collected by centrifugation at 16,000 x g at 4°C for 10min and pellets washed twice in ice cold acetone and then resuspended din 2X SDS-laemmli buffer and proteins separated on 1 10% SDS-PAGE gel.

### Co-Immunoprecipitation

100×106 PAS 835 (Fkh2:13XMYC; Clr6:TAP) or PAS 837 (Fkh2:13XMYC; Sds3:TAP) cells were grown, lysed and the lysate clarified as above (sucrose gradient analysis) but in the following lysis buffer: 20mM HEPES pH7.6, 150mM NaCl, 1mM EDTA, 0.5% IGEPAL NP-40, 10% glycerol, 1mM PMSF, 200μM phenantroline, pepstatin A, leupeptin, aprotinin and 1X of EDTA-free protease-inhibitor cocktail (Roche)). 500μL total protein was incubated with 20μL IgG-Sepharose 6 resin (GE healthcare) for 2 hrs at 4°C with rotation. Beads were washed 4 times with lysis buffer with 350mM NaCl instead of 150mM. Proteins were eluted off washed beads in 20μL 2X SDS-laemmli buffer, separated on SDS-PAGE gels and blotted as below.

### Western blot analysis

Proteins were transferred to low-fluorescence PVDF membranes (Bio-rad) at 90min at 200mA at 4°C. Membranes were blocked in 1:1 mixture of 1XPBS : Intercept PBS blocking buffer (LiCor) and then incubated with anti-PAP (Sigma, P1291, lot 92557) or anti-MYC (Biolegend, 626802, lot B274036) at 1:1,000 either overnight at 4°C or 90 min at RT. Membranes were washed 4X in the presence of 0.2% Tween-20 and then incubated with fluorescent anti-mouse (800nm, Rockland, 610-145-003, lot 34206) and anti-rabbit (680nm, Cell Signaling, 5366P, lot 9) at 1:5,000 and 1:15,000 respectively for 1hr at RT. Membranes were washed 4X as above, transferred to 1X PBS and imaged on a LiCor Odyssey CLx imager.

### Data Availability

Raw and processed ChIP-seq data will be available via GEO upon publication of the manuscript. Code for analysis of ChIP-seq data will be deposited in Zenodo (10.5281/zenodo.5122951). Software version information will be included in conda environment yml files (2021_seqTools.yml for command line data processing and 2021_Renv.yml for analysis via R/Bioconductor).

## Acknowledgments

We thank the Michael N Boddy lab for their generous gift of strains expressing tagged Clr6 Iʺ complex subunits. We thank Sy E Redding, Carol A Gross, Douglas Myers-Turnbull and Kamir Hiam for helpful discussions on data acquisition, analysis, and interpretation, Arthur Molines for help with microscopy experiments, and Sandra Catania and Hiten D Madhani for support for chromatin immunoprecipitation experiments. This work was supported by grants from the National Institutes of Health (DP2GM123484), the National Science Foundation (NSF 2113319), and the UCSF Program for Breakthrough Biomedical Research (partially funded by the Sandler Foundation) to BA-S, and the ARCS Foundation Scholarship and Hooper Graduate Fellowship to RAG. HN and CT were supported by National Science Foundation Graduate Research Fellowships (grant number 1650113). This work was supported by grants awarded to SB from the European Union Network of Excellence EpiGeneSys (HEALTH-2010-257082). SB is a Heisenberg Program Fellow (BR 3511/BR 5-1) and Member of the Collaborative Research Center 1064 (Project-ID 213249687) funded by the Deutsche Forschungsgemeinschaft (DFG, German Research Foundation) and acknowledges infrastructural support. Flow cytometry and FACS data were generated in the UCSF Parnassus Flow Cytometry Core which is supported by the Diabetes Research Center (DRC) grants NIH P30 DK063720. The library preparation and sequencing were carried out at the DNA Technologies and Expression Analysis Cores at the UC Davis Genome Center, supported by NIH Shared Instrumentation Grant 1S10OD010786-01.

---

**Figure 1 Supplement 1:**
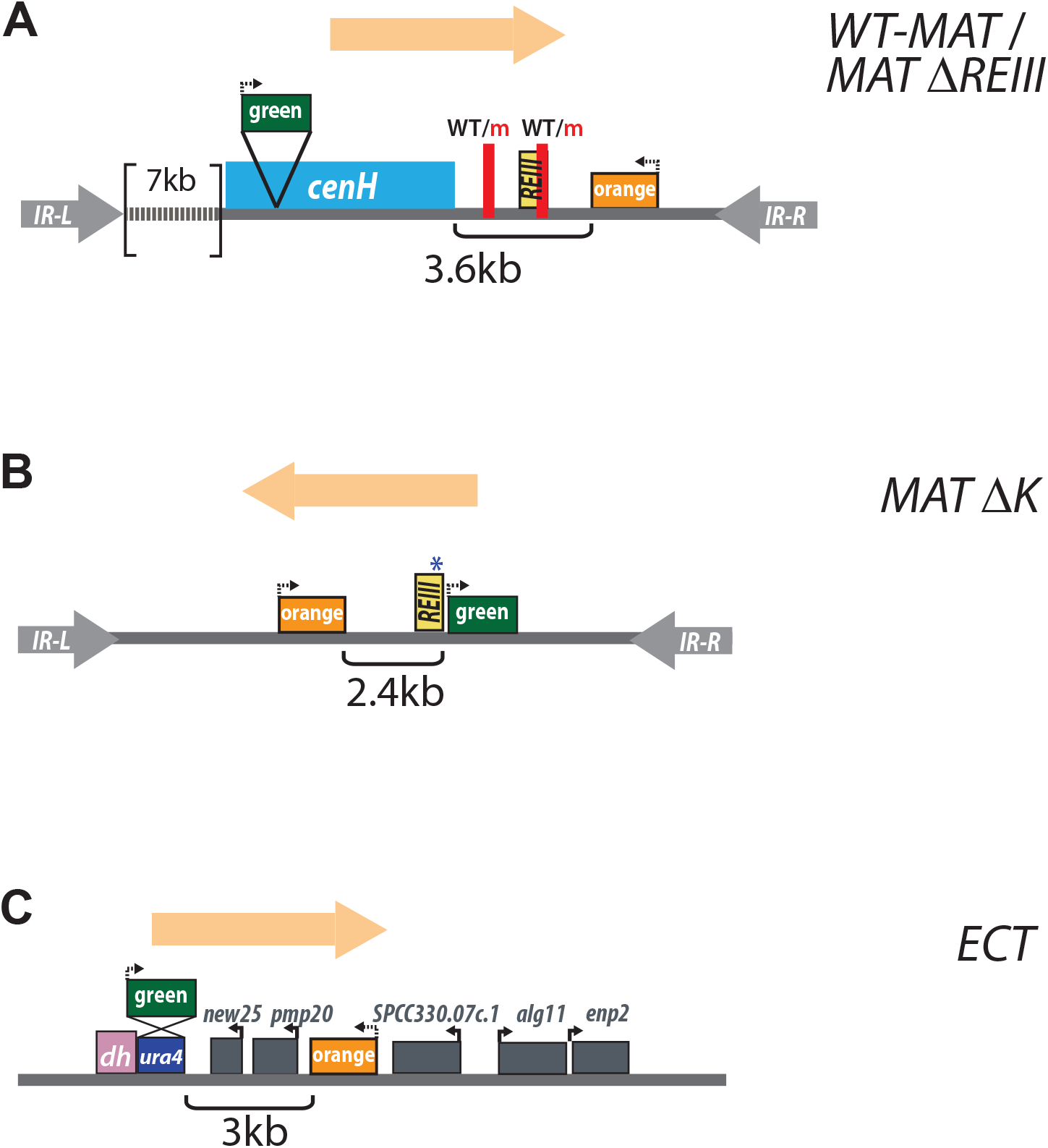
Screen chromatin contexts with “green” and “orange” reporters. To-scale diagrams of the heterochromatin spreading sensors (without the euchromatically placed “red” reporter) in the 4 chromatin contexts used for the spreading screen (as in Greenstein et al 2018). The direction in which spreading is analyzed (“green” to “orange”) is indicated per chromatin context. **A.** *WT MAT* and *MAT ΔREIII.* These two contexts are similar, except that *MAT ΔREIII* contains two short 7bp deletions of the two Atf1/Pcr1 DNA binding sites near *REIII*, inactivating it. The first binding site is not included in *REIII*, per the definitions of (Thon et al. 1999) and (Jia et al. 2004). **B.** *MAT ΔcenH*. **C.** *ECT*. In Greenstein et al 2018, the distance between “green” and “orange” was varied for B. and C. contexts, without changes to the qualitative behavior of spreading.

**Figure 1 Supplement 2:**
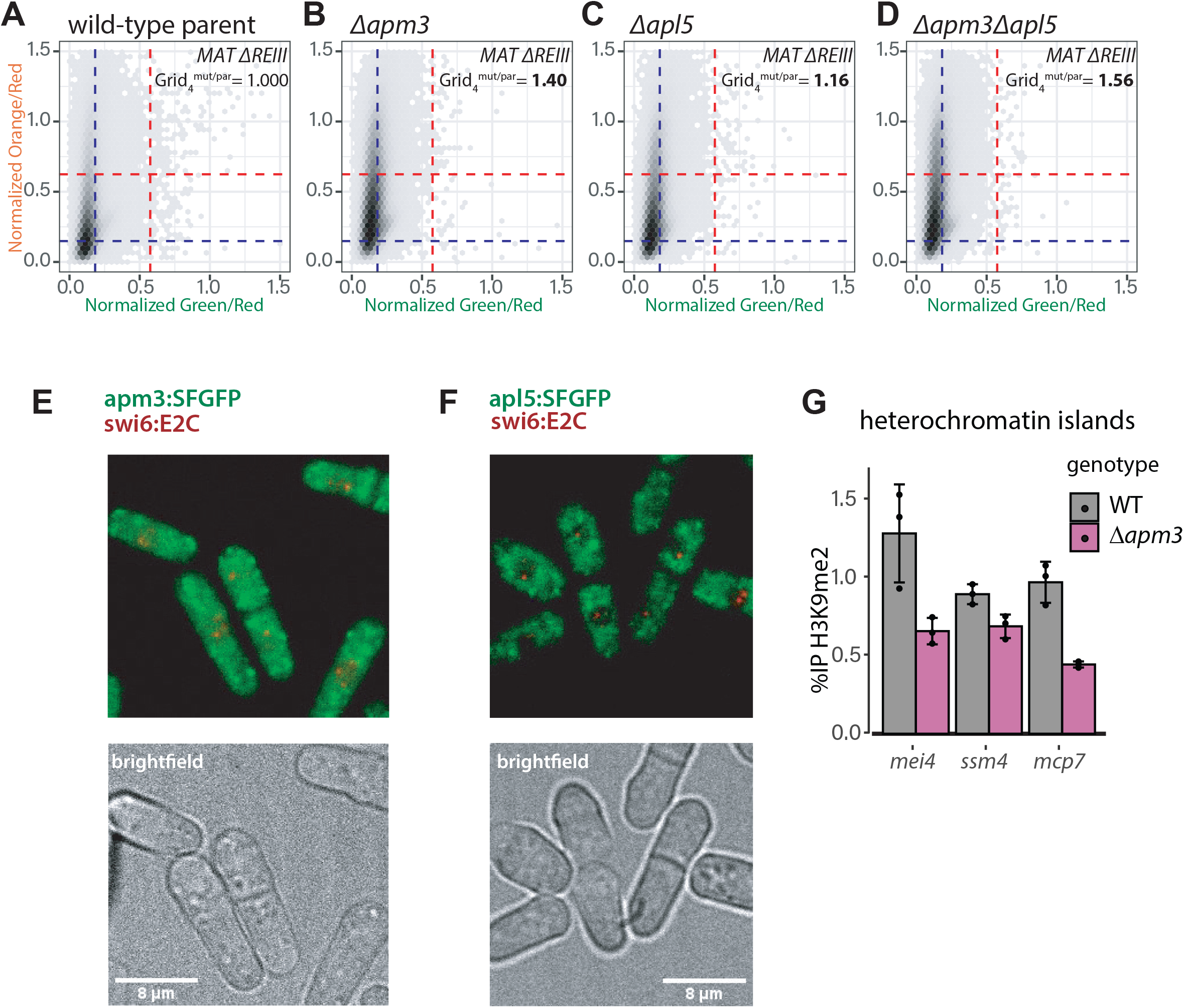
*apm3* and *apl5*, coding for nuclear-cytosolic and cytosolic proteins, respectively, act together in modulation of heterochromatin spreading. **A.-D.** 2D density hexbin plots of *de novo* generated *Δapm3* (B.), *Δapl5* (C.), and *Δapm3Δapl5* double mutant (D.) compared to the wild-type *MAT ΔREIII* parent (A.). The Fold change of Grid_4_^mut/par^ is indicated in the plot. At least 3 independent isolates of each genotype are combined in each plot. **E.** Apm3:SFGFP is distributed in the cytosol and nucleus. Apm3:SFGFP was expressed from its native locus and co-expressed with Swi6:E2C. Swi6:E2C labels nuclear heterochromatin. Z-projection overlays of the Apm3:SFGFP and Swi6:E2C on top, and a brightfield image on the bottom. **F.** Apl5:SFGFP is largely nuclear excluded. Apl5:SFGFP was expressed from its native locus and co-expressed with Swi6:E2C. Swi6:E2C labels nuclear heterochromatin. Z-projection overlays of the Apl5:SFGFP and Swi6:E2C on top, and a brightfield image on the bottom. **G.** *Δapm3* exhibits a mild defect in H3K9me2 accumulation at heterochromatin islands. H3K9me2 ChIP-qPCR in wild-type parent *MAT ΔREIII* or *Δapm3* mutant.

**Figure 1 Supplement 3:**
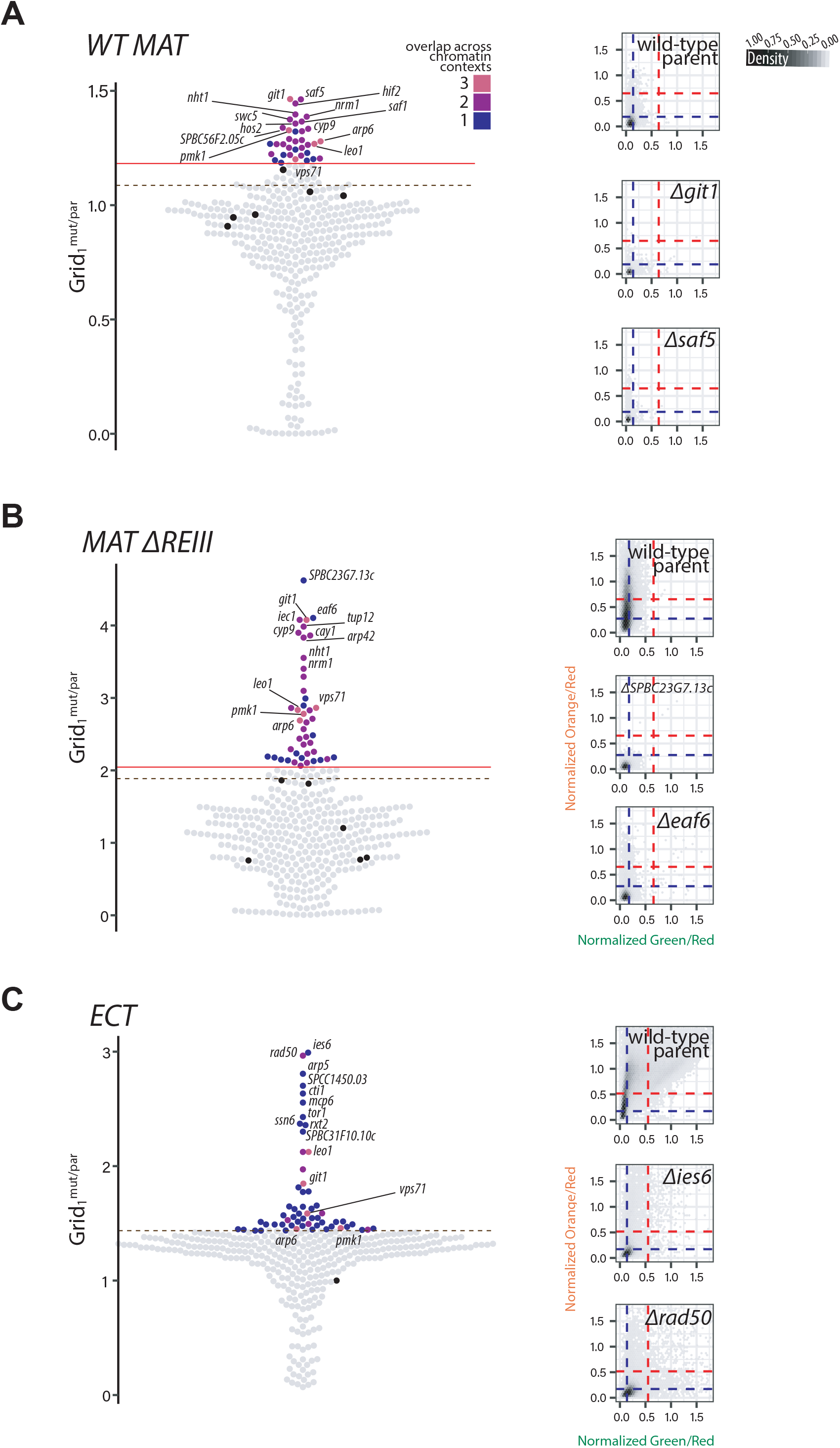
Swarm- and 2D density hexbin plots for gain of spreading. **A.** LEFT: Beeswarm plots of Grid_1_ for *WT MAT* gain of spreading hits. The top 10 hits are all annotated, and below those hits, mutants that show overlap with 2 other chromatin contexts are additionally annotated. Red line, 2SD above the Grid_1_ of wild-type parent isolates(black dots); dashed brown line, the 85^th^ percentile; Dot color, number of chromatin contexts with loss of spreading phenotype over the cutoff. RIGHT: *WT-MAT* 2D-density hexbin plots of the wild-type parent, and the two top gain of spreading hit of this chromatin context. Dashed blue lines indicate the values for repressed fluorescence state and dashed red lines indicate values for fully expressed fluorescence state. **B.** As in A. but for *MAT ΔREIII*. **C.** As in A. but for *ECT*.

**Figure 1 Supplement 4:**
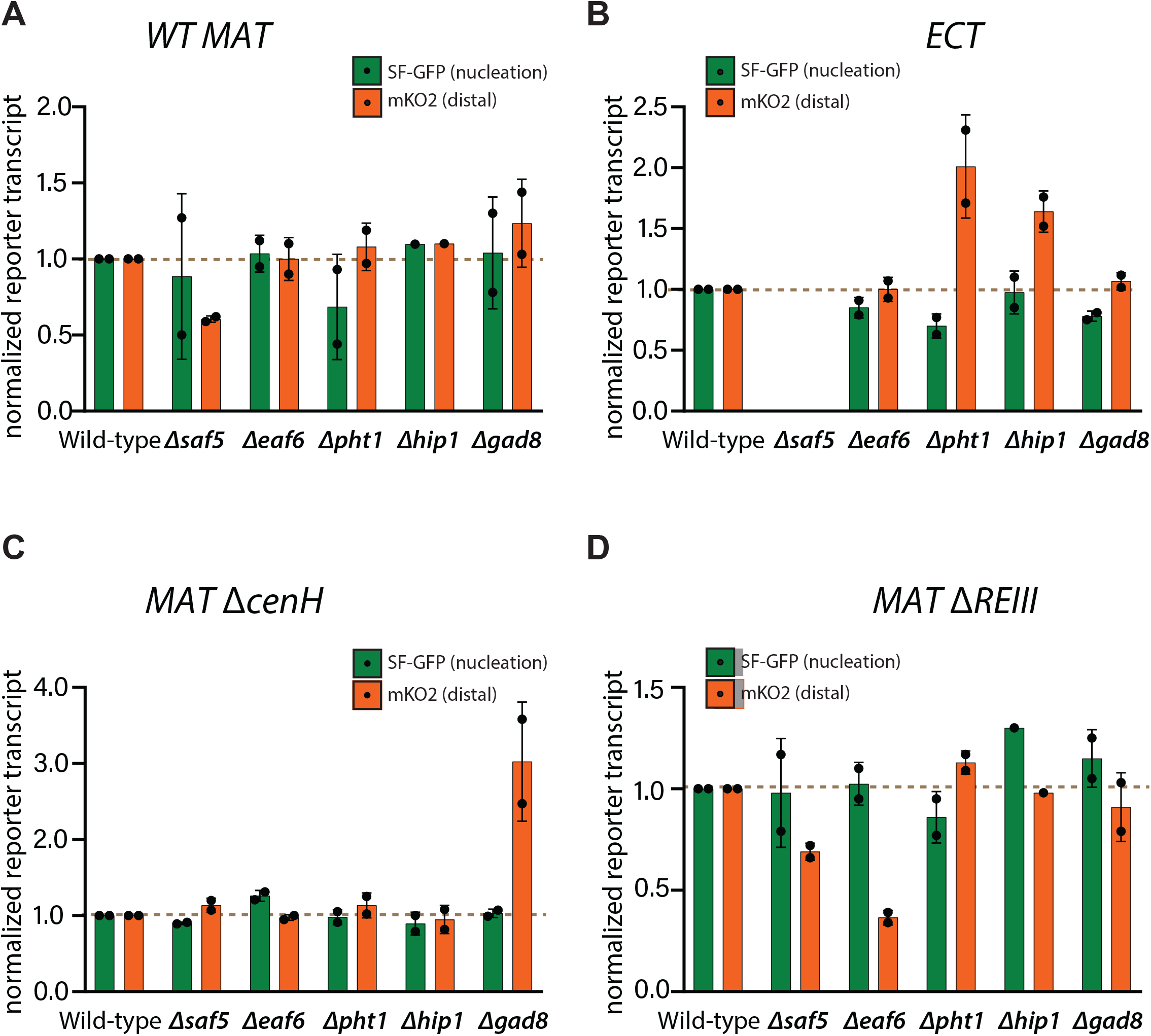
RT-qPCR validations of selected chromatin-context unique loss and gain of spreading hits. 5 moderate- to strong hits in the loss and gain of spreading category that are partially or fully chromatin context-specific were selected for validations: *saf5* (gain of spreading in *WT MAT*, and moderately in *MAT ΔREIII*), *eaf6* (gain of spreading only in *MAT ΔREIII*), *pht1* and *hip1* (loss of spreading only in *ECT*), and *gad8* (strong loss of spreading in *MAT ΔcenH* and mildly in *ECT*). RT-qPCRs for SF-GFP (“*green*”-nucleation) and mKO2 (*“orange”,* spreading) transcripts normalized to the *act1* transcript are shown for **A.** *WT-MAT,* **B.** *ECT,* **C.** *MAT ΔcenH,* and **D.** *MAT ΔREIII.* Error bars indicate 1SD of 2 biological replicates generated independently from the screen. Note we could not recover *Δsaf5* mutants in *ECT*.

**Figure 1 Supplement 5:**
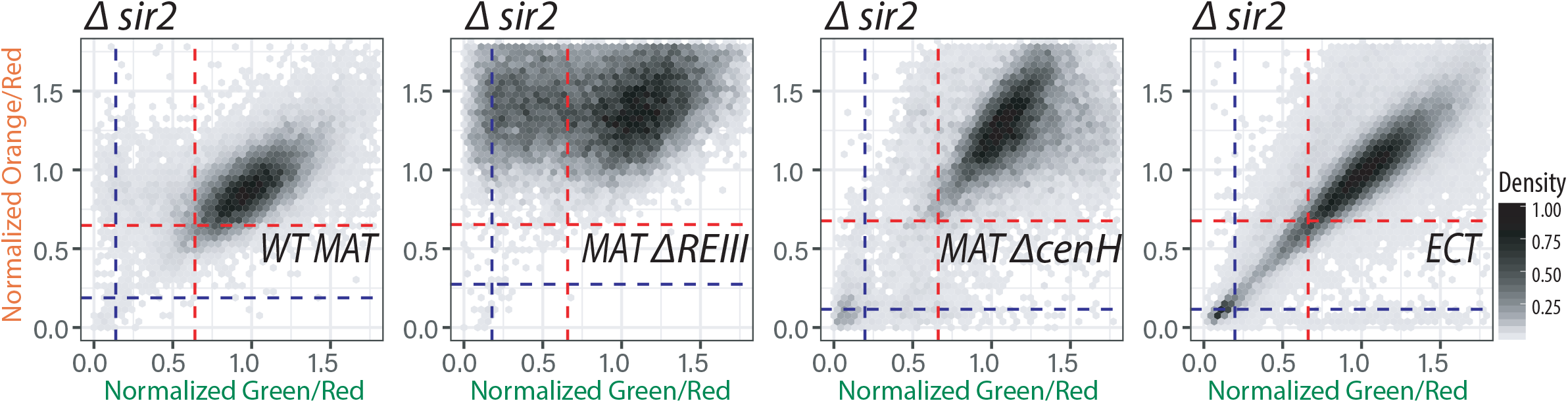
Class III HDAC family Sir2 is required for heterochromatin silencing. 2D density hexbin plots of *Δsir2* mutants in each chromatin context from the screen. Mutation in *sir2* causes a loss of silencing phenotype in all examined chromatin context.

**Figure 2 Supplement 1:**
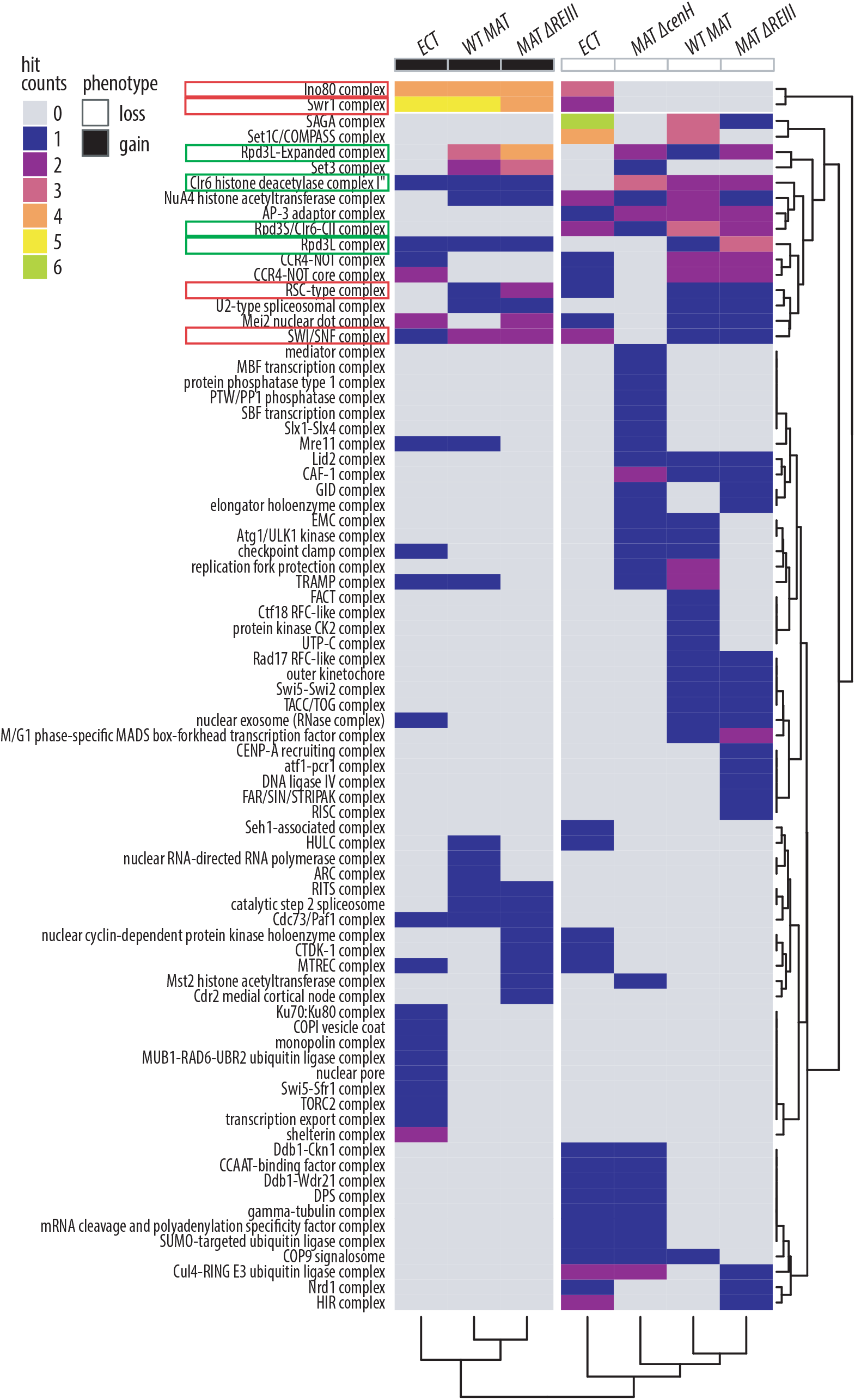
Heterochromatin spreading is regulated by sets of unique and common protein complexes across different chromatin contexts. Heatmap of GO complex annotations for hits in each category and strain. Rows, representing GO complexes annotated to genes within the screen that were identified as hits, are arranged via hierarchical clustering. Columns are defined by the hit phenotype (loss of spreading – white; gain of spreading – black), and each screen chromatin context is indicated at the top. The columns were clustered by hierarchical clustering and the dendrogram was cut to define 2 branches. Red boxes, chromatin remodeling complexes; Green boxes, Clr6 complexes (Note that Rpd3L Expanded includes Set3C).

**Figure 2 Supplement 2:**
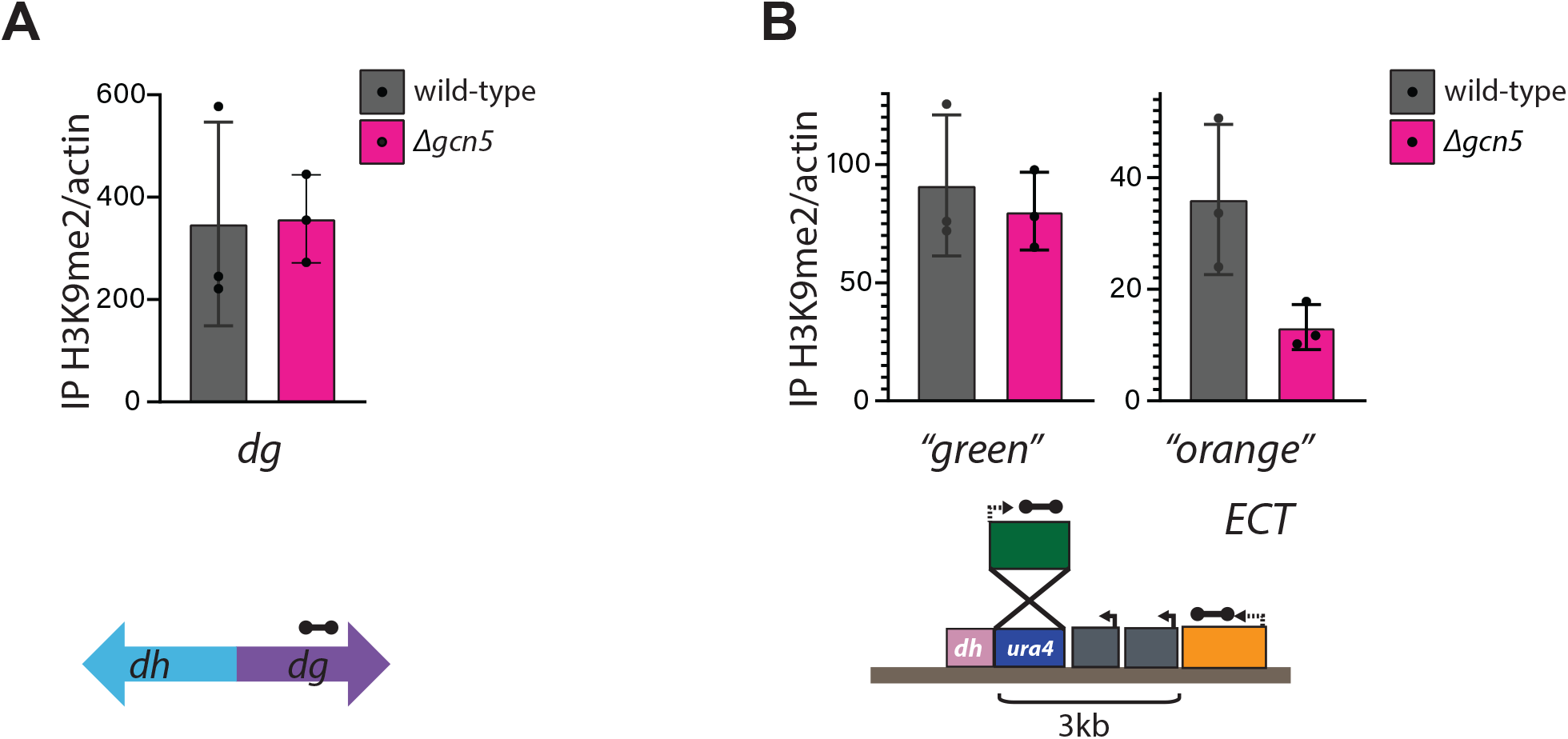
*gcn5* is specifically required for H3K9me2 spreading at the *ECT* heterochromatin spreading sensor, but not pericentromeric heterochromatin. *act1-*normalized ChIP-qPCR for H3K9me2 in *ECT* wild-type parent or the *de novo* generated *Δgcn5* mutant at **A.** the pericentomeric *dg* element, and **B.** the heterochromatin spreading sensor at the *ura4* locus in *ECT.* Dumbbells indicate qPCR amplicons. Error bars indicate 1SD of 3 biological replicates.

**Figure 2 Supplement 3:**
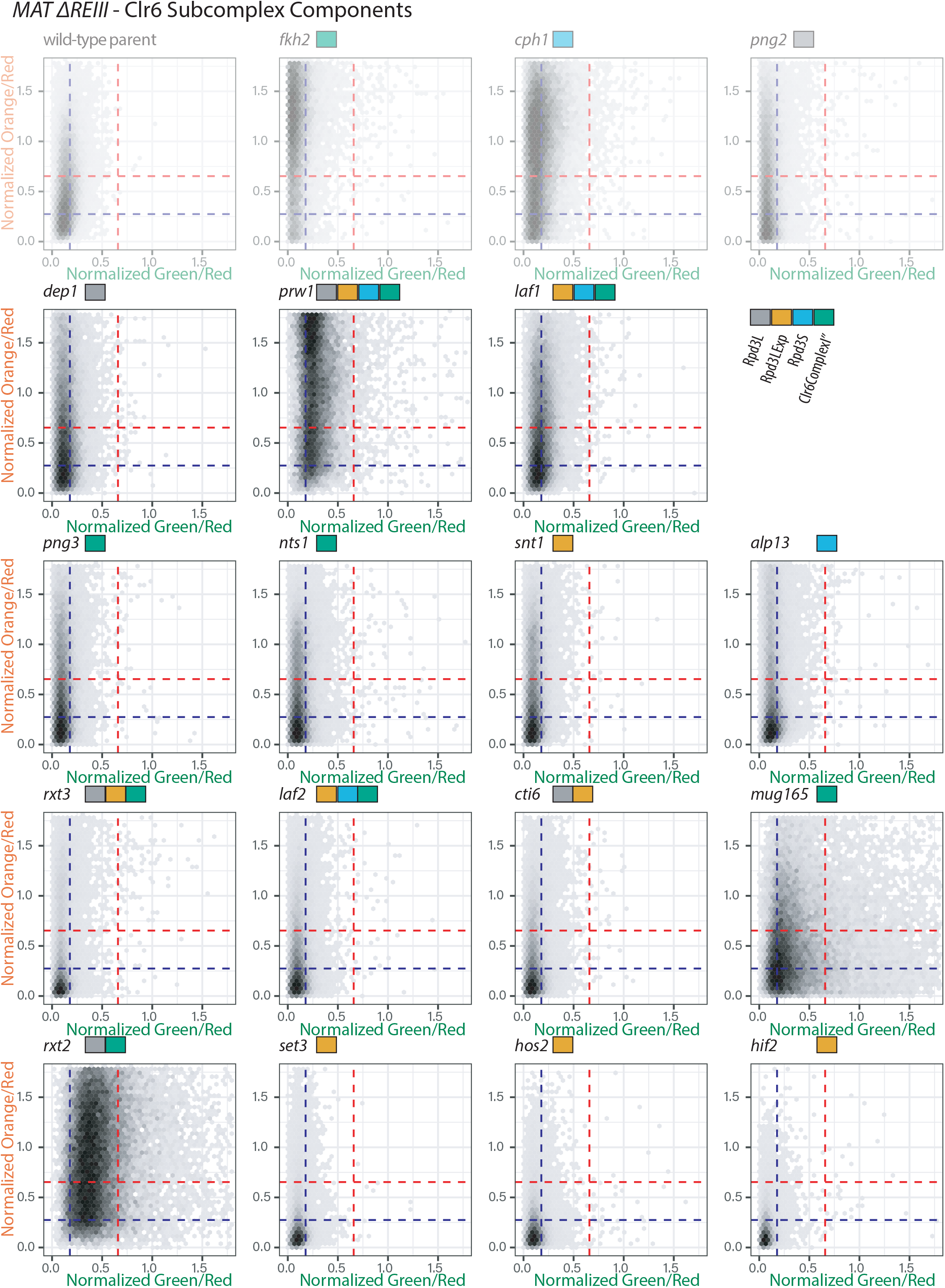
2D density hexbin plots for all Clr6 complex subunit screen mutants in *MAT ΔREIII*. 2D density hexbin plots of all Clr6 complexes gene mutants from the screen, corresponding to Figure 2C in *MAT ΔREIII* context. The mutants are arranged in descending order of Grid_3+_ ^mut/par^; in *MAT ΔREIII* only *Δfkh2, Δcph1, Δpng2, Δdep1, Δprw1* and *Δlaf1* were identified as loss of spreading phenotype. Original *MAT ΔREIII* wild type parent and mutants shown in Figure 2 are shown here again (with transparency) for comparison. GO complex annotations are indicated next to each mutant by colored boxes.

**Figure 3 Supplement 1:**
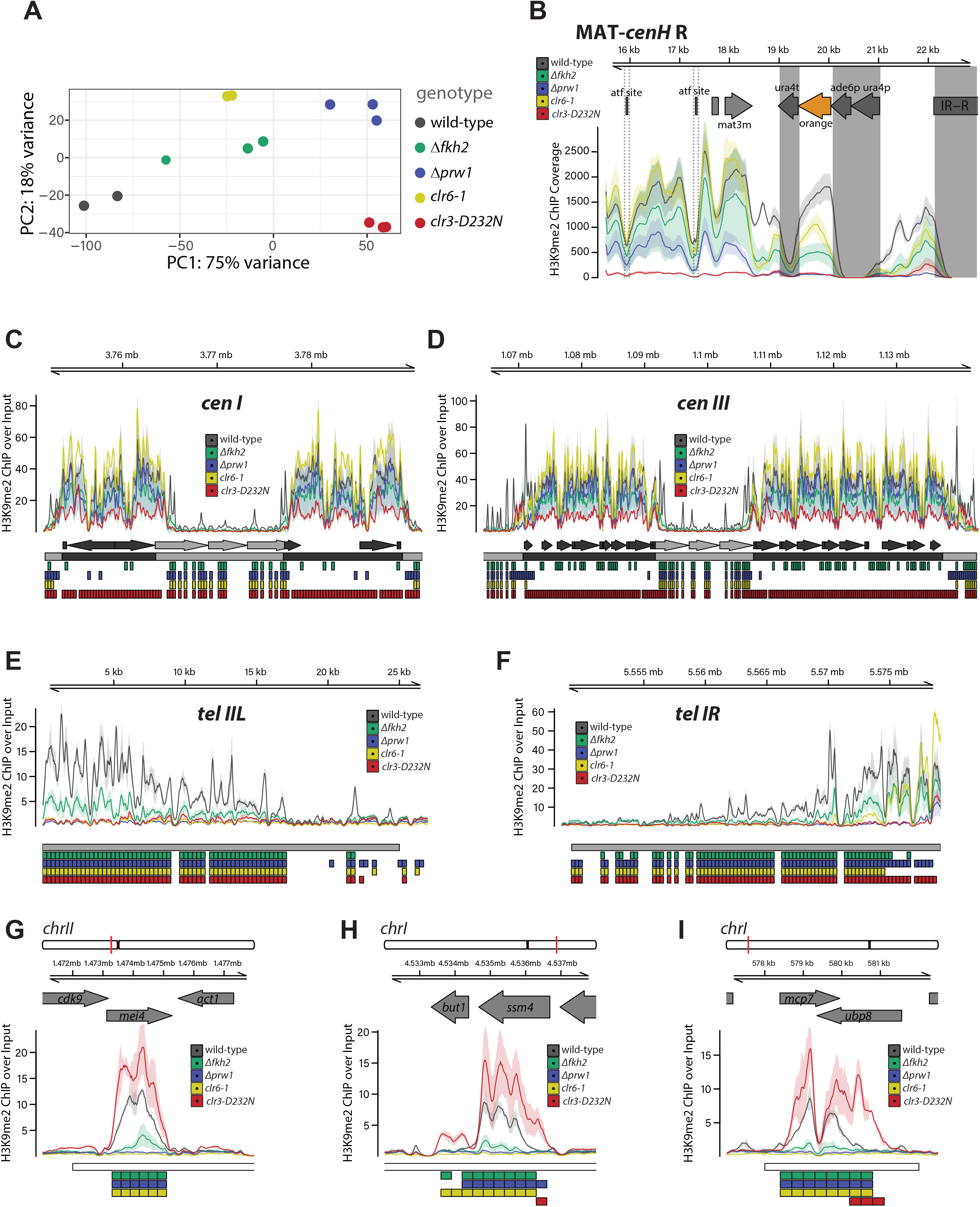
Fkh2-containing Clr6 complexes direct H3K9me2 spreading at multiple genomic regions. **A.** Principal Component Analysis was performed on the normalized counts in the top 500 most variable bins passing a threshold for global enrichment of H3K9me2 signal (see methods). The first two principal component values are plotted for each sample with genotypes as defined in the legend. **B.** Alignment was performed to a custom mating type region contig in *MAT ΔREIII* with green and orange color cassettes. ChIP signal for 10bp intervals is plotted as mean (line) and 95% confidence interval (shade) for each genotype (see legend). ChIP signal represents the coverage of each interval adjusted for the sequencing depth of the full genome bam file relative to that of the full genome bam file with the lowest depth. Features of interest are annotated above the signal tracks. During data processing for alignment to this custom contig, reads mapping to multiple locations within the reference sequence were removed. For this reason, there is little to no signal over regions that are homologous within this reference including *ura4p*/*ade6p* at the color cassette promoters, *ura4t* at the color cassette terminators, and *IR-L* and *IR-R* elements (these regions are shaded). Signals at feature “atf site” (indicated by dashed box) are reduced as both 7bp sites in *MAT ΔREIII* are deleted. **C.-F.** Signal tracks plots for indicated centromeres and telomeres as in the main text. No nucleator sequences are present on subtelomeres I right and II left so the first annotation row below the signal tracks is empty. **G.-I.** Signal tracks for *mcp7, mei4, ssm4* islands are plotted as in the main text. Genes are annotated above the signal tracks. Below the signal tracks the following annotations are present in order from top to bottom: (1) previously identified euchromatin embedded H3K9me2 heterochromatin region (“island”, “HOOD”, or “region”) annotated as a white box. (2-5) 300bp regions determined to be significantly differentially enriched for the comparisons between *Δfkh2* and wild-type [green], *Δprw1* and wild-type [blue], *clr6-1* and wild-type [yellow], *clr3-D232N* and wild-type [red] respectively annotated as colored boxes.

**Figure 3 Supplement 2:**
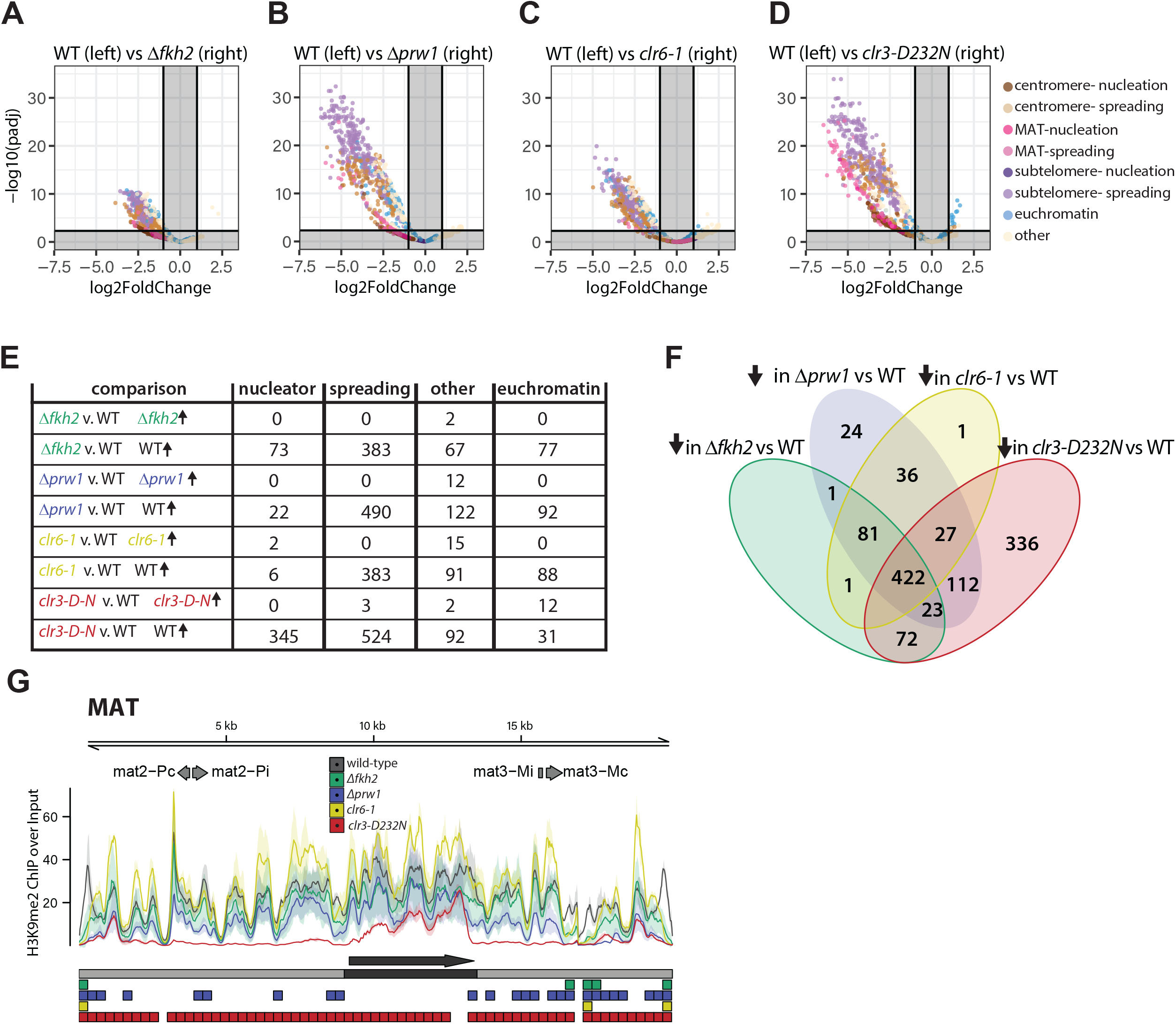
Fkh2-containing Clr6 complexes contribute primarily to H3K9me2 spreading, while Clr3 is required for H3K9me2 accumulation at all heterochromatin regions except islands. **A.-D.** Volcano plots were generated as in Figure 3. Dots are colored by their annotation to nucleation or spreading zones broken down by heterochromatin location (pericentromere, subtelomere, MAT) or presence within a previously identified euchromatin embedded H3K9me2 heterochromatin region. **E.** The number of regions called as significant in each direction for each of the pairwise comparisons is tabulated per each category of genomic feature. **F.** The overlap of regions identified as significantly reduced in H3K9me2 signal in each mutant vs WT is compared in a Venn Diagram. **G.** Signal tracks for H3K9me2 ChIP-Seq in *MAT ΔREIII* at the full mating_type_region contig from the Pombase reference genome, as in Figure 3E. Gene annotations for the mating type cassette genes are additionally included above the signal tracks. Note that most Clr6 subunits only show defects in spreading to the right of *cenH,* indicated with a dark grey arrow.

**Figure 3 Supplement 3:**
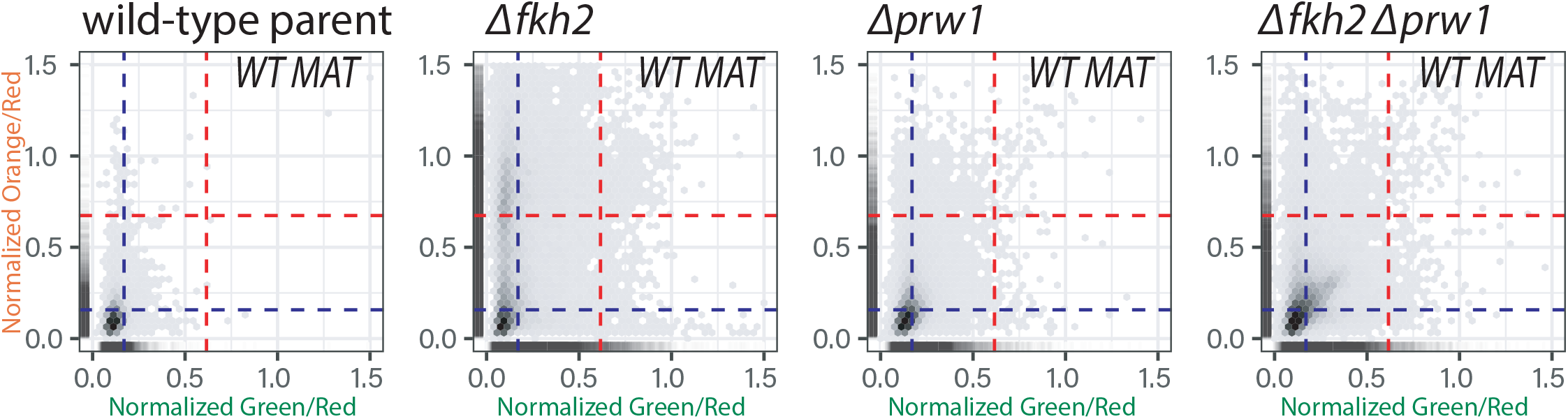
Validation of *Δfkh2, Δprw1 and Δfkh2 Δprw1* double mutant spreading defects in *WT MAT*. *Δfkh2, Δprw1 and Δfkh2 Δprw1* mutants were re-created *de novo* in *WT MAT.* 2D density hexbin and rug plots of indicated backgrounds are shown.

**Figure 3 Supplement 4:**
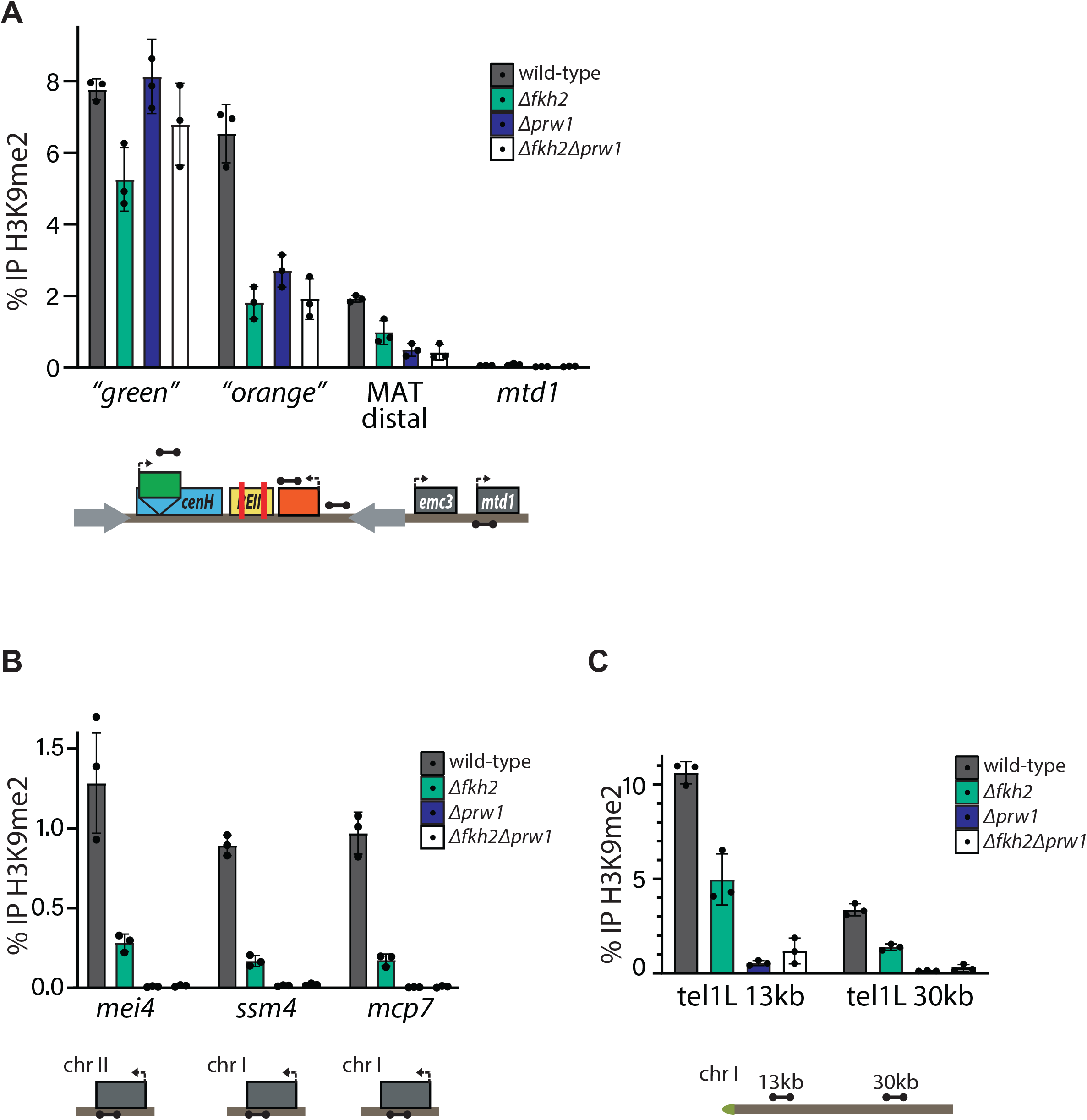
Fkh2 and Prw1 act together in spreading H3K9me2. **A.** H3K9me2 ChIP -qPCR at the MAT locus in wild-type *MAT ΔREIII, Δfkh2, Δprw1,* and the *Δfkh2Δprw1* double mutant. **B.** As in A., at indicated heterochromatin islands. **C.** As in A., at *tel IL*. Error bars represent 1SD of 3 biological replicates.

**Figure 4 Supplement 1:**
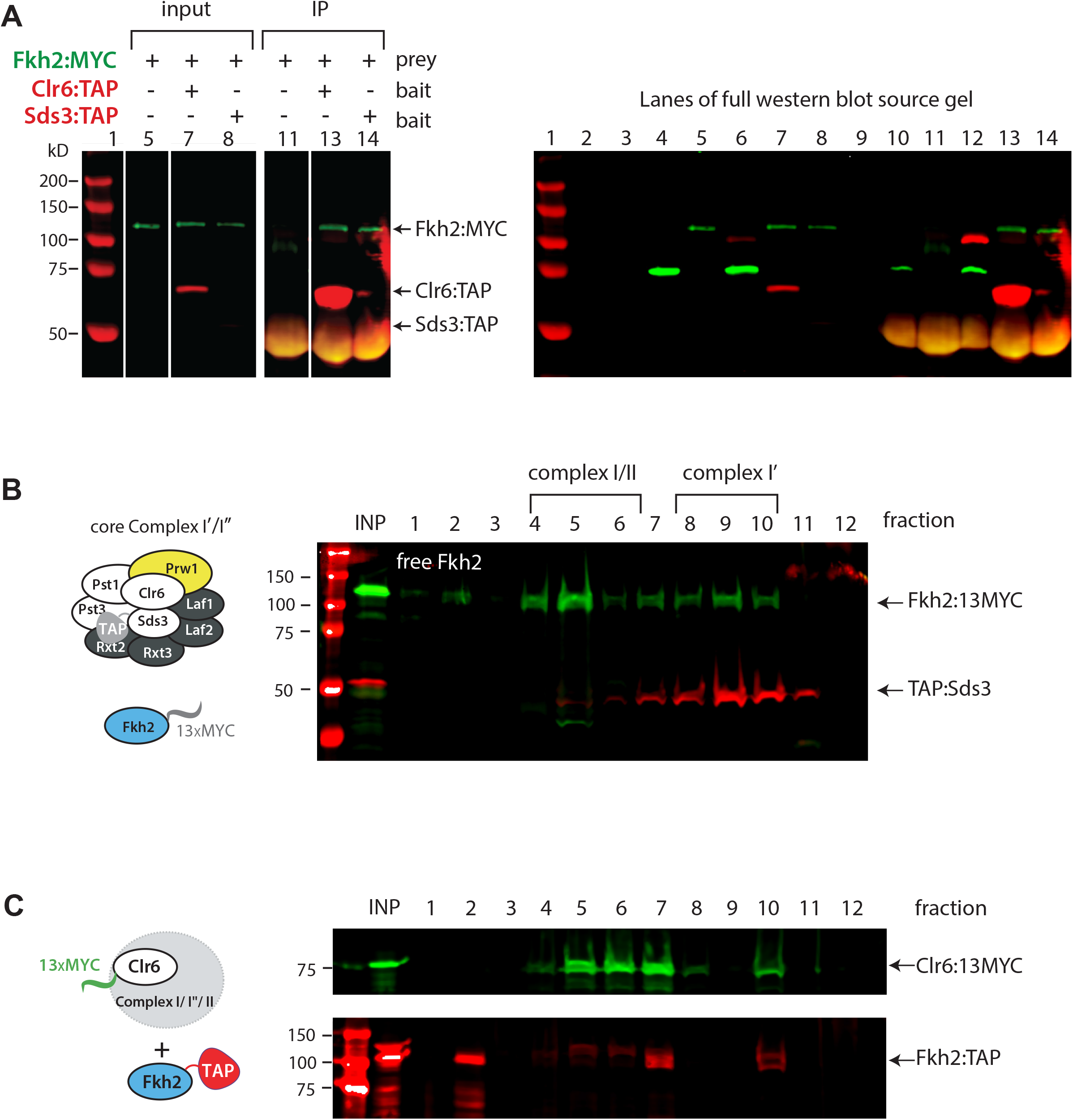
Fkh2 is a constituent member of Clr6 complexes. **A.** Co-Immunoprecipitation experiment with baits Clr6-TAP or Sds3-TAP and prey Fkh2-MYC. LEFT: Western blot against indicated proteins for the Co-IP experiment. RIGHT shows the entire western blot, including lanes unrelated to the co-IP experiment (2-4,5,10,12). **B.** Sucrose density gradient for whole cell extracts of cells containing Sds3-TAP, a signature of complex I/Iʺ, and Fkh2:MYC. Gradient and Western as in Figure 4. **C.** Single channel Western blots of Figure 4 A.

**Figure 4 Supplement 2:**
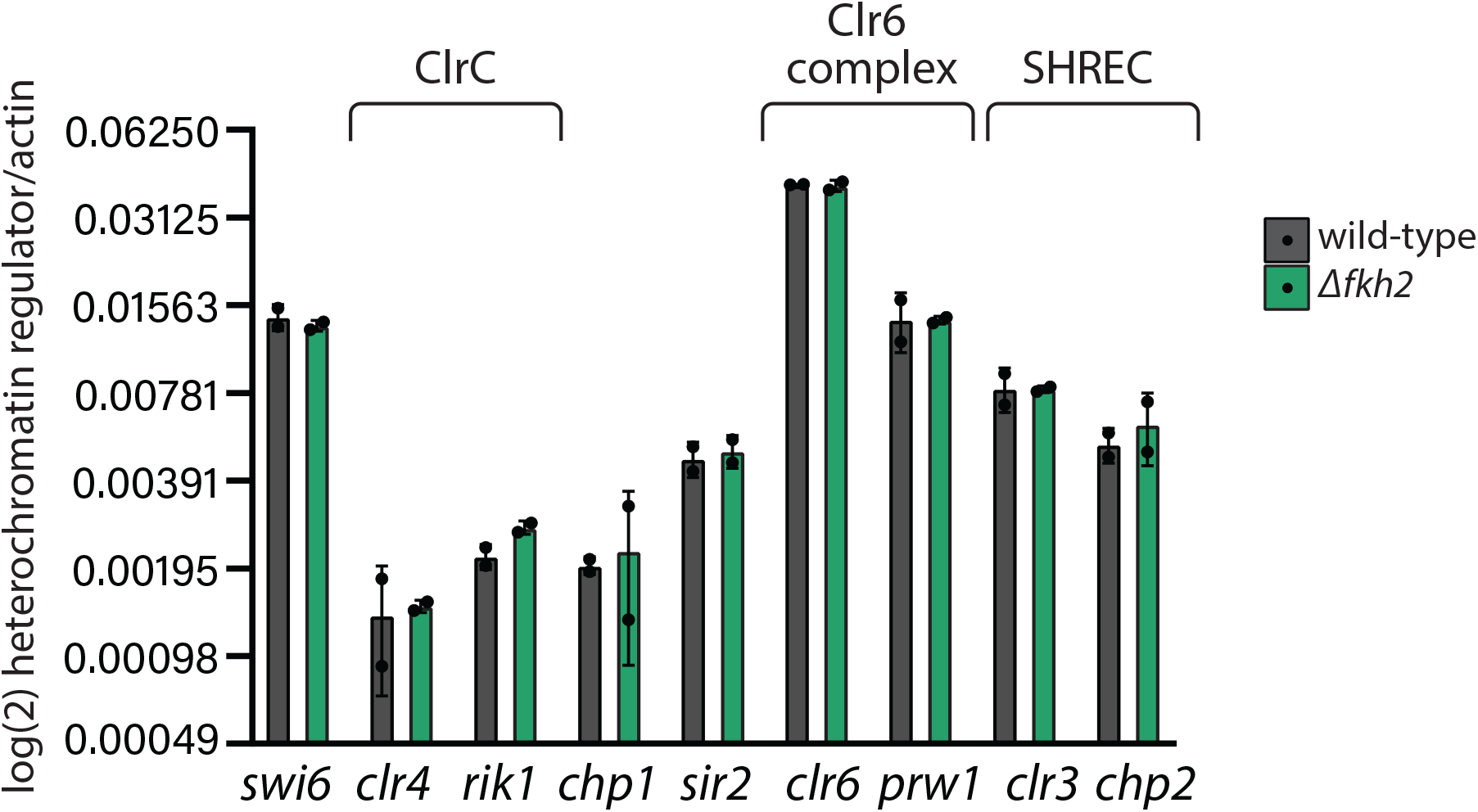
Fkh2 does not affect transcription of core heterochromatin regulators. RT-qPCR of indicated core heterochromatin regulators, including representatives of ClrC, Clr6, and SHREC in *MAT ΔREIII* wild-type or *Δfkh2* cells. Heterochromatin regulator transcripts are normalized to the *act1* transcript and shown on a log2 scale, given the wide distribution of transcript abundance between indicated regulators. Error bars indicate 1SD of 2 biological replicates.

